# Crystallographic fragment binding studies of the *Mycobacterium tuberculosis* trifunctional enzyme suggest binding pockets for the tails of the acyl-CoA substrates at its active sites and a potential substrate channeling path between them

**DOI:** 10.1101/2024.01.11.575214

**Authors:** Subhadra Dalwani, Alexander Metz, Franziska U. Huschmann, Manfred S. Weiss, Rik K. Wierenga, Rajaram Venkatesan

**Affiliations:** Faculty of Biochemistry and Molecular Medicine, University of Oulu, Oulu, Finland; Department of Pharmaceutical Chemistry, Philipps-University Marburg, Marburg, Germany; Macromolecular Crystallography, Helmholtz-Zentrum Berlin, Berlin, Germany

**Keywords:** CoA-dependent enzymes, electrostatic surface, lipid metabolism, substrate channeling, tuberculosis, fatty acid β-oxidation, trifunctional enzyme

## Abstract

The *Mycobacterium tuberculosis* trifunctional enzyme (MtTFE) is an α_2_β_2_ tetrameric enzyme in which the α-chain harbors the 2*E*-enoyl-CoA hydratase (ECH) and 3*S*-hydroxyacyl-CoA dehydrogenase (HAD) active sites, and the β-chain provides the 3-ketoacyl-CoA thiolase (KAT) active site. Linear, medium, and long chain 2*E*-enoyl-CoA molecules are the preferred substrates of MtTFE. Previous crystallographic binding and modelling studies have identified binding sites for the acyl-CoA substrates at the three active sites as well as the NAD^+^ binding pocket at the HAD active site. These studies have also identified three additional CoA binding sites on the surface of MtTFE that are different from the active sites. It has been proposed that one of these additional sites could be of functional relevance for substrate channeling (by surface crawling) of reaction intermediates between the three active sites. Here, in a crystallographic fragment binding study with MtTFE crystals 226 fragments were screened, resulting in the structures of 17 MtTFE-fragment complexes. Analysis of the 143 fragment binding events shows that the ECH active site is the ‘binding hotspot’ for the tested fragments, with 50 binding events. The mode of binding of the fragments bound at the active sites provides additional insight on how the long chain acyl moiety of the substrates can be accommodated at their proposed binding pockets. In addition, the 24 fragment binding events between the active sites identify potential transient binding sites of reaction intermediates relevant for possible channeling of substrates between these active sites. These results provide a basis for further studies to understand the functional relevance of these binding sites and to identify substrates for which channeling is crucial.

**Synopsis:** Crystallographic fragment binding studies of the *Mycobacterium tuberculosis* trifunctional enzyme (MtTFE) have resulted in 143 binding events of 17 fragments out of 226 investigated fragments, suggesting functional sites with respect to substrate binding and substrate channeling.

## Introduction

A large portion of the genome of *Mycobacterium tuberculosis (Mtb)* codes for enzymes involved in lipid metabolism (Cole *et al*., 1998). Our understanding of the importance of lipid metabolism at the various stages of infection by *Mtb* has improved over the years and several enzymes involved in the pathways of lipid metabolism are possible drug targets (Mi *et al*., 2022). It has been shown that the genes that code for fatty acid metabolism in general as well as specifically for the β-oxidation pathway are upregulated during the intracellular stages of infection (Rohde *et al*., 2012; Schnappinger *et al*., 2003). *Mtb* is capable of switching its metabolic preference during the latent stage of infection in order to utilize host-derived fatty acids as a source of carbon rather than glucose or glycerol (Wilburn *et al*., 2018). *Mtb* has several copies of genes coding for the enzymes of the β-oxidation cycle; however, only a single trifunctional enzyme (MtTFE) that catalyzes the last three out of four reactions of the β-oxidation cycle (**Fig. 1**), is present in *Mtb*, in contrast to other bacteria such as *Escherichia coli (E. coli)* which has two TFEs (Sah-Teli *et al*., 2023, 2019). MtTFE has been characterized to be an α_2_β_2_ heterotetrameric multifunctional enzyme complex (**Fig. 1**) encoded by the *fadA* (β subunit) and *fadB* (α subunit) genes (Venkatesan & Wierenga, 2013).

**Figure 1.**
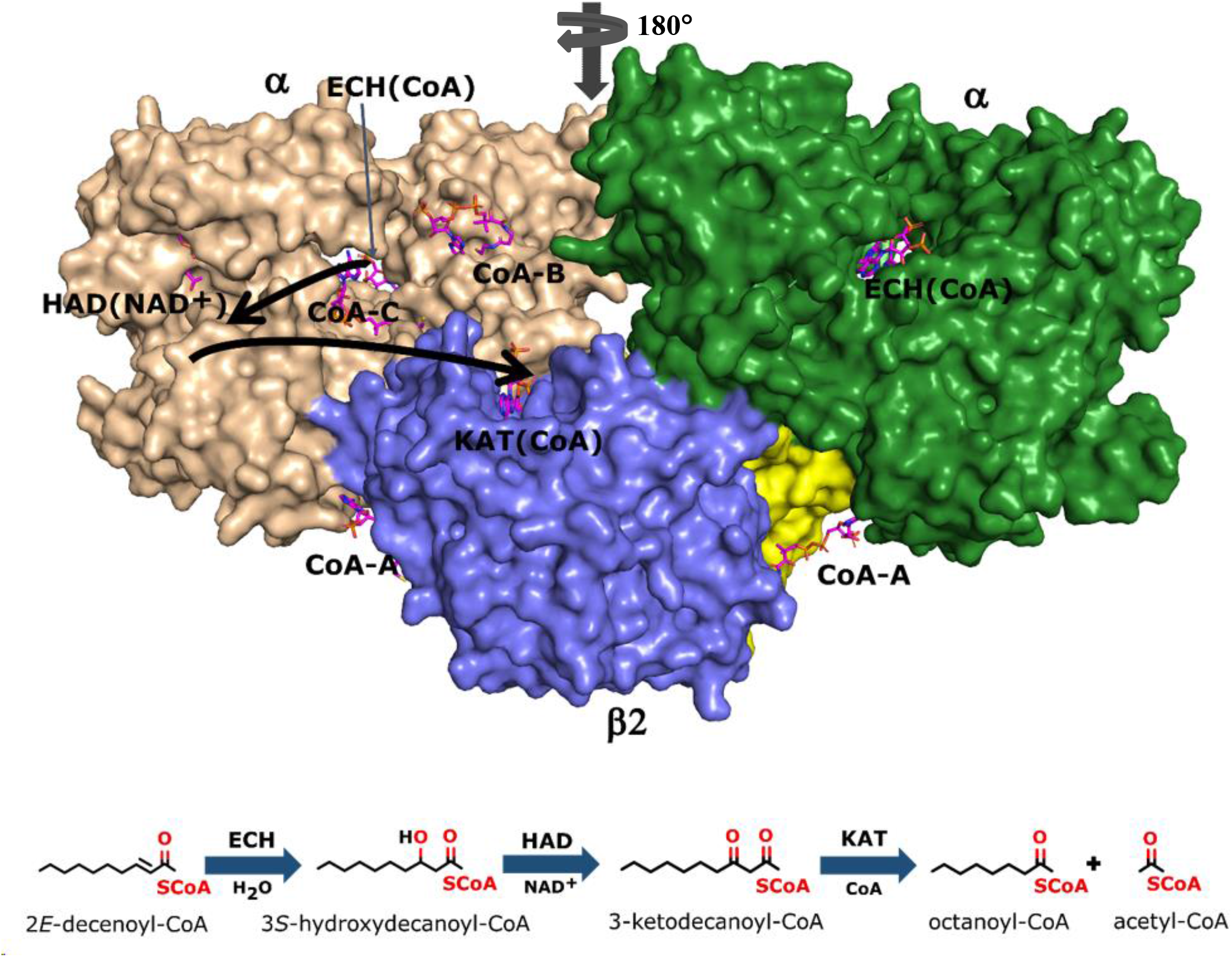
The TFE tetramer with its ECH, HAD and KAT active sites. (*upper panel*) The two α-chains (wheat and green) are assembled on top of the β_2_ thiolase dimer (cyan, yellow). The vertical arrow visualises the two-fold axis of the α_2_β_2_-tetramer. The ECH, HAD and KAT active sites labeled as ECH(CoA) (with thin arrow), HAD(NAD^+^) and KAT(CoA). CoAs bound at the additional CoA binding sites are labeled as CoA-A, CoA-B and CoA-C. The curved thick arrows identify the path between the ECH and HAD active site and between the HAD and KAT active site. (*lower panel*) Schematic representation of the reactions catalyzed by the ECH, HAD and KAT active sites.

Bioinformatics studies have suggested that four TFE subfamilies exist: (i) the mammalian mitochondrial TFE, (ii) the mycobacterial TFE (MtTFE), (iii) the bacterial aerobic TFE of *E*.*coli* (EcTFE) and *Pseudomonas fragi* (PfTFE) and (iv) the bacterial anaerobic TFE (of *E. coli*, anEcTFE) (Venkatesan & Wierenga, 2013). Experimental evidence for the presence of substrate channeling by the multifunctional enzymes of the β-oxidation pathway for specific substrates exists for the rat peroxisomal multifunctional enzyme (type 1) (RnMFE1) (Yang *et al*., 1986), EcTFE (Yang *et al*., 1986), mitochondrial TFE from pig heart (Yao & Schulz, 1996) and mitochondrial TFE from humans (Nada *et al*., 1995). While biochemical assays provide kinetic evidence for the existence of substrate channeling in a given multifunctional enzyme system, structural studies are essential to provide insight into a possible mechanism and for identifying the transient binding sites. Substrate channeling offers several advantages in cellular metabolism (Sweetlove & Fernie, 2018). The mechanism of substrate channeling is dependent on the nature of the substrate. For small molecules it concerns substrate tunneling through the matrix of the protein, for example in the well-studied bifunctional enzyme tryptophan synthase (Barends *et al*., 2008), and for large polar molecules it concerns surface crawling (over bulk solvent exposed surface area) as observed for the bifunctional enzyme thymidylate synthase-dihydrofolate reductase (TS-DHFR) (Anderson, 2017). In some cases, partial channeling of substrates has been observed (Baker *et al*., 2012).

Each of the β-oxidation TFEs form α_2_β_2_ heterotetramers (M=240kDa) in which the β_2_ dimer forms the core of the tetramer, however, their quaternary assemblies differ, due to different modes of assembly of the α chains onto the β_2_ dimer (Ishikawa *et al*., 2004; Liang *et al*., 2018; Sah-Teli *et al*., 2023, 2020; Venkatesan & Wierenga, 2013; Xia *et al*., 2019). MtTFE, EcTFE and PfTFE are soluble enzymes whereas HsTFE and anEcTFE are membrane associated. The different quaternary assemblies of the four TFE subfamilies, the electrostatic surface features between the active sites, as well as membrane association have been speculated to play a role in the possible mechanisms for substrate channeling between the active sites of these TFEs (Ishikawa *et al*., 2004; Sah-Teli *et al*., 2023, 2020). The reaction intermediates of the three TFE reactions are always negatively charged due to the phosphate groups of the CoA moiety and it has been proposed that positively charged residues on the surfaces of these proteins are used to guide the negatively charged intermediates between the three active sites. In addition to the charged surfaces, also the formation of reaction chambers, due to the mode of assembly, has been proposed to play a role in substrate channeling. For HsTFE and anEcTFE membrane association is also hypothesized to be important in this respect (Sah-Teli *et al*., 2023; Xia *et al*., 2019).

The crystal structure of MtTFE in its unliganded and CoA bound forms identified the binding sites for its CoA-conjugated substrates at each of its three active sites (CoA(ECH), CoA(HAD) and CoA(KAT)) (Venkatesan & Wierenga, 2013) as well as the binding site for the NAD^+^ co-factor at the HAD active site (Dalwani *et al*., 2021). A structural analysis of MtTFE highlighted the presence of positively charged residues between the ECH and HAD active sites and a neutral surface path between the HAD and KAT active sites (Dalwani *et al*., 2021; Venkatesan & Wierenga, 2013). These crystallographic binding studies revealed also three additional CoA binding sites on the surface of MtTFE, referred to as the CoA-B(ECH2), CoA-A(HAD/KAT) and CoA-C(ECH/HAD) sites (Dalwani *et al*., 2021) (**Fig. 1**).

Crystallographic fragment screening has been established as an attractive and useful tool to identify regions on the surface of proteins to which small molecule ligands (M<250Da) bind with weak affinity (Martin & Noble, 2022; Wollenhaupt *et al*., 2020). These studies have shown that functional binding sites can be identified by fragment binding studies (Carbery *et al*., 2022; Radeva *et al*., 2016), even if the affinity for the fragments is low and difficult to detect with other biophysical methods (Price *et al*., 2017; Schiebel *et al*., 2016). Further studies are required to establish the key features of the binding determinants at these surface patches (Anderson, 2022; Czub *et al*., 2022; Davies *et al*., 2011; Hilario *et al*., 2016). The information obtained from a crystallographic fragment binding study has also been used in the early stages of drug discovery research, in particular by combining the structural information from multiple fragment hits for generating larger lead compounds with higher affinity, which subsequently has led to the development of new potential drug molecules (Boby *et al*., 2023; Erlanson *et al*., 2016; Heightman *et al*., 2018). Via this approach also new functional binding sites have been identified (Krojer *et al*., 2020; Shumilin *et al*., 2012; Skaist Mehlman *et al*., 2023). Here, fragment screening has been used as a tool to identify weak transient binding sites on the surface of MtTFE. The possible functional importance of these sites is discussed in the context of the mode of binding of the acyl-tails of the acyl-CoA substrates at the three active sites and in the context of substrate channeling of reaction intermediates between the three active sites.

## Methods

### Protein expression and purification

MtTFE was expressed in *E. coli* BL21 (DE3) and purified using a previously standardized protocol (Venkatesan & Wierenga, 2013). Once purified, protein was concentrated in storage buffer (20 mM 4-(2-hydroxyethyl)-1-piperazineethanesulfonic acid (HEPES) pH 7.2, 120 mM KCl, 1mM EDTA and 1 mM NaN_3_), flash frozen using liquid nitrogen and stored at -70 °C for further use.

### Protein crystallization

MtTFE was crystallized in its unliganded form using a protocol as described previously (Venkatesan & Wierenga, 2013). For this, 0.5 μL of MtTFE solution (6 mg mL^-1^ in storage buffer), was mixed with the crystallization well solution containing 2 M (NH_4_)_2_SO_4_ in 100 mM tris(hydroxymethyl)aminomethane (Tris) buffer at pH 8.5 in a 1:1 ratio using the Mosquito nanodispensing robot (TTP Labtech) and crystallized by vapor diffusion in hanging drops at room temperature. The plates were imaged using the Formulatrix Rock imager (RI54) at regular time intervals and the formation of the crystals was monitored using the IceBear software (Daniel *et al*., 2021).

### Choice of fragment libraries and preparing the fragment containing drops

#### (a) Library from Helmholtz-Zentrum Berlin (HZB)

96 fragments selected for general crystallographic fragment screening purposes based on size, diversity, presence as ligand in other PDB entries, cost and availability (Huschmann *et al*., 2016) were pre-spotted in a single fragment per spot format as provided by HZB. 100 nL of MtTFE crystallization solution (2 M (NH_4_)_2_SO_4,_ 100 mM Tris pH 8.5) was pipetted onto each deposited fragment spot, such that completely soluble fragments would be at a final concentration of 100 mM in the fragment drop. Subsequently, 50 μL of the same MtTFE crystallization solution was immediately pipetted as reservoir solution into the wells of all the spotted fragments, the plate was sealed and fragment drops were allowed to equilibrate against the crystallization solution for 24 hours at room temperature before starting the crystal soaking experiment.

#### (b) Compounds from Philipps-University Marburg

In order to mimic the negatively charged acyl-CoA substrates/intermediates of MtTFE, 130 negatively charged small molecule compounds were provided as pre-weighed powder in tubes or as aqueous solution. A 1M, 0.5 M or 0.25M stock concentration of each of the provided compounds was made by dissolving the powder in either 100% DMSO or 50% DMSO/water or used as the provided aqueous solution, as specified in **Table S1**. Not all the compounds dissolved completely, when the calculated volume of solvent was added. In those cases, the suspension of the compound was vortexed vigorously before being used for spotting. For the spotting, respectively 100, 200 or 400 nL of each fragment suspension was deposited onto a TTP Labtech plate as a single fragment per deposited spot. The drops were dried off at room temperature or in an incubator at 25 °C for up to 48 hours until no solvent was visible. Once the solvent had completely evaporated, 70 μL of MtTFE crystallization solution (2M (NH_4_)_2_SO_4_, 100 mM Tris, pH 8.5) was pipetted into the wells of the TTP Labtech plate and the spotted ligands were reconstituted with 500 nL of the same well solution using the Mosquito nano dispensing robot, such that the nominal concentration of each of the compounds was 200 mM. The drops were sealed and subsequently allowed to equilibrate for 24 hours at room temperature before starting the crystal soaking experiment.

### Fragment soaking

For fragment soaking, the individual soaking experimental approach was adopted where 5-10 unliganded crystals of MtTFE were transferred into each of the 226 pre-spotted fragment containing drops and incubated at room temperature for at least 24 hours before cryo-cooling in liquid nitrogen. The suitability of the MtTFE crystals for the fragment soaking experiments was assessed by visual inspection of images obtained from IceBear (Daniel *et al*., 2021). Mostly larger crystals (larger than 100 μm in diameter) that had grown in drops with a single or a few crystals per drop were selected. Once selected, crystals were harvested and manually transferred by using a loop into the sitting drop that contained the pre-spotted fragments. Subsequently, the crystals were allowed to equilibrate in the fragment solution for 24-48 hours at room temperature, directly cryo-cooled in liquid nitrogen and subsequently stored in liquid nitrogen for the diffraction experiment. In some cases, crystals cracked or dissolved during the soaking step before they could be frozen and in some cases, the crystals that survived the soaking step diffracted to less than 3.2 Å resolution. Thus, these could not be checked further for binding studies and were excluded from the data processing and analysis steps.

### Data collection, data processing and structure refinement

Frozen fragment-soaked crystals were transferred in dry shippers to various European synchrotrons for data collection. X ray diffraction data sets were collected at different beamlines at BESSY II, DLS, MAX IV and PETRA III. All data collection was done at a temperature of 100 K. Data reduction was either performed manually using XDS (Kabsch, 2010) and AIMLESS (Evans & Murshudov, 2013) or by using the various data processing pipelines, available at the different synchrotron facilities including XDSAPP (Sparta *et al*., 2016; Krug *et al*., 2012), xia2 (Winter, 2010), xia2_3dii (Winter, 2010), fast_dp (Winter & McAuley, 2011), xia2_DIALS (Winter *et al*., 2018), autoPROC (Vonrhein *et al*., 2011) and STARANSIO (Tickle, I.J. *et al*., 2018). Only data sets that could be processed to a resolution of 3.2 Å or better were used for molecular replacement calculations for obtaining initial phases. The data collection and data processing statistics are listed in **Table S2** and **Table S3**. The model used as search model in molecular replacement was derived from PDB ID 4B3H, after removing bound ligands and water molecules, using either PHASER (McCoy *et al*., 2007) or MOLREP (Vagin & Teplyakov, 2010). Positioned coordinates after molecular replacement were used as the initial model to perform iterative rounds of model building and refinement using COOT (Emsley *et al*., 2010), phenix.refine of Phenix (Afonine *et al*., 2012; Liebschner *et al*., 2019) and REFMAC5 of CCP4 (Murshudov et al., 2011; Potterton et al., 2018; Winn et al., 2011). Ligands were built in their electron density only after several rounds of manual model building and refinement and after adding water molecules. The structure quality was assessed using MolProbity (Williams *et al*., 2018) as well as by inspecting the validation report from the PDB validation server (Feng *et al*., 2021; Smart *et al*., 2018*b*).

The quality of fit between ligand model and observed electron density was assessed both by manually checking the electron density maps as well as by using the real-space correlation coefficient (RSCC) of the bound fragment (as provided in the PDB validation report) (Smart *et al*., 2018*a*). An RSCC value of 0.8 was chosen to filter bound fragments with a good fit to density (∼93% of the fragment binding events had an RSCC value of 0.8 or higher), however in a few cases, ligands with lower RSCC values were also kept for analysis (∼6% with RSCC between 0.76 and 0.79 and there are 2 ligands with RSCC values between 0.76 and 0.72) (**Table S4**). The relatively liberal cut-off value for the RSCC values for some of the fragments (for those with RSCC values between 0.7 and 0.8) was selected keeping in mind the size of the complex and resolution of the data sets and the fragments with lower RSCC values were retained either based on evidence of binding at the same site in the other copy of the tetramer, or because of the binding of another similar fragment at the same binding site. Occupancies were not refined but from the information of the difference maps in some cases the occupancies were manually set (**Table S4)**. The final refinement statistics are listed in **Table S2** and **Table S3**. A summary of the data processing and refinement statistics of the refined structures is provided in **Table 1**.

**Table 1.**
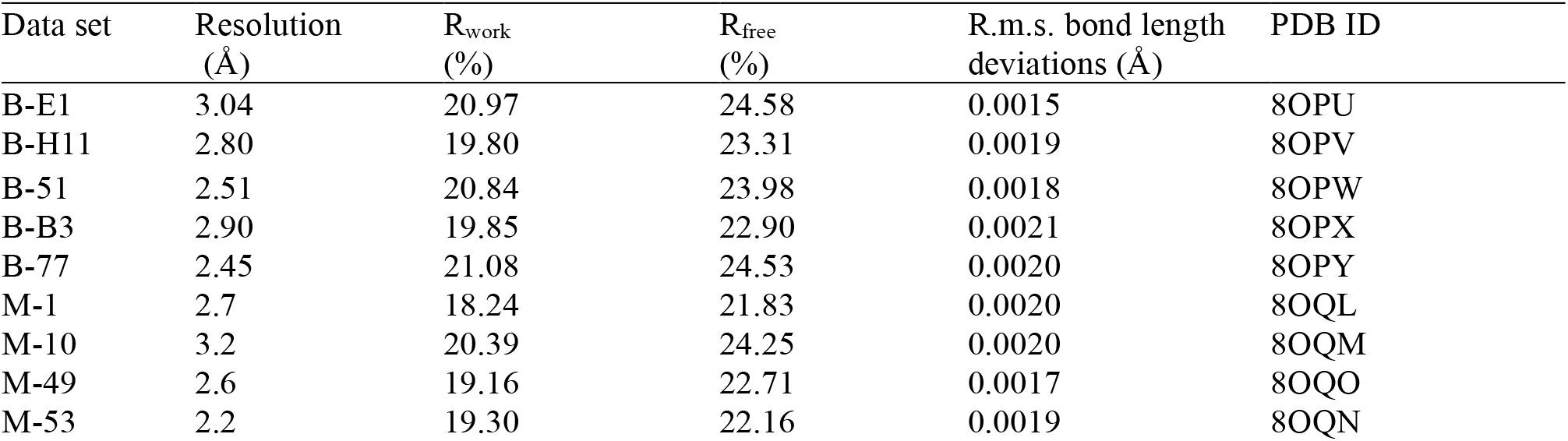
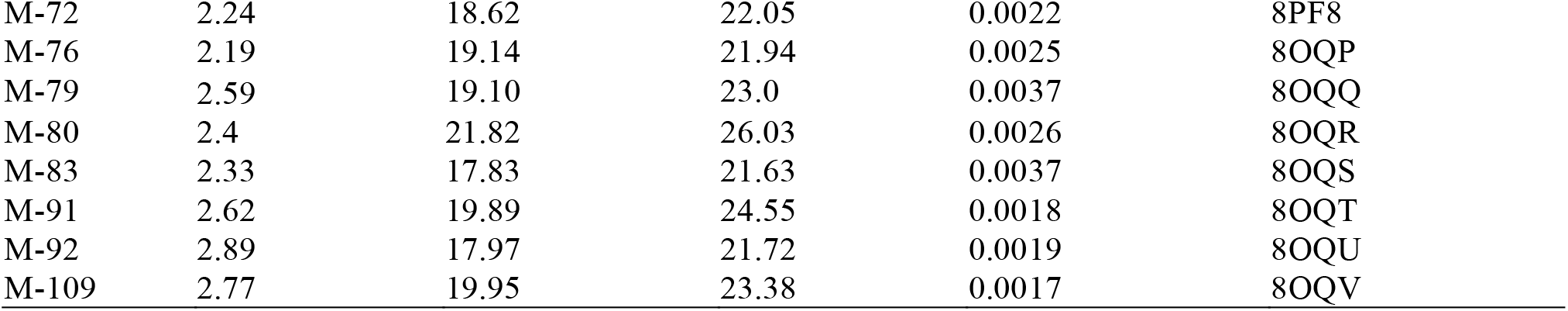
Summary of the refinement statistics of the structures of the 17 MtTFE-fragment complexes. Detailed data collection and refinement statistics are provided in **Table S2** and **Table S3**. The covalent structures of the fragments are provided in **Table S1**.

| Data set | Resolution<br>(Å) | R<sub>work</sub><br>(%) | R<sub>free</sub><br>(%) | R.m.s. bond length<br>deviations (Å) | PDB ID |
| --- | --- | --- | --- | --- | --- |
| B-E1 | 3.04 | 20.97 | 24.58 | 0.0015 | 8OPU |
| B-H11 | 2.80 | 19.80 | 23.31 | 0.0019 | 8OPV |
| B-51 | 2.51 | 20.84 | 23.98 | 0.0018 | 8OPW |
| B-B3 | 2.90 | 19.85 | 22.90 | 0.0021 | 8OPX |
| B-77 | 2.45 | 21.08 | 24.53 | 0.0020 | 8OPY |
| M-1 | 2.7 | 18.24 | 21.83 | 0.0020 | 8OQL |
| M-10 | 3.2 | 20.39 | 24.25 | 0.0020 | 8OQM |
| M-49 | 2.6 | 19.16 | 22.71 | 0.0017 | 8OQO |
| M-53 | 2.2 | 19.30 | 22.16 | 0.0019 | 8OQN |
| M-72 | 2.24 | 18.62 | 22.05 | 0.0022 | 8PF8 |
| M-76 | 2.19 | 19.14 | 21.94 | 0.0025 | 8OQP |
| M-79 | 2.59 | 19.10 | 23.0 | 0.0037 | 8OQQ |
| M-80 | 2.4 | 21.82 | 26.03 | 0.0026 | 8OQR |
| M-83 | 2.33 | 17.83 | 21.63 | 0.0037 | 8OQS |
| M-91 | 2.62 | 19.89 | 24.55 | 0.0018 | 8OQT |
| M-92 | 2.89 | 17.97 | 21.72 | 0.0019 | 8OQU |
| M-109 | 2.77 | 19.95 | 23.38 | 0.0017 | 8OQV |

### Ligand restraints

The restraints for the fragments were generated according to the structural formulas as provided in **Table S1**. Coordinates and geometry restraints for the HZB compounds were provided by HZB. Coordinates and restraints for the set of compounds that were provided by Philipps-University Marburg were generated from SMILES strings using the GRADE websever (Smart *et al*.). For fragment M-1 (PDB Chemical Component ID A9J), REEL (Moriarty *et al*., 2017) of phenix was used to fix the ligand geometry in the restraints file available from the PDB. For fragment M-72 the electron density maps showed the mode of binding of 15 fragments. In 8 binding events it concerned molecules having the structure as provided in **Table S1** (PDB Chemical Component ID JXL). In 2 cases a dimeric derivative (**Fig. S1**) was observed (PDB Chemical Component ID YLN) (covalently bound to the side chain of His(−9), of the A and B chains), and in 5 other binding events either the monomeric form (4 binding events) or the dimeric form (1 binding event) were suggested by the electron density map to be modified by a glycerol molecule (PDB Chemical Component ID YLZ and PDB Chemical Component ID YMK, respectively). The geometry of these modified fragments are in agreement with the boron chemistry (Diaz & Yudin, 2017). The restraints of the covalent link between the dimer fragment and the histidine side chain were generated using the program JLigand of CCP4 (Nicholls *et al*., 2021).

### Mass spectrometry of solubilized fragments

Fragments M-80, M-83, M-91, M-92 and M-109 were provided as aqueous solutions created through hydrolysis of the corresponding sulfonyl chlorides by dissolving them in 1M aqueous NaOH. The conversion of these sulfonyl chlorides to the respective sulfonic acids (**Table S1**), was checked using electrospray ionization mass spectrometry (ESI-MS). In addition, the oligomeric state of fragment M-72 was also checked using ESI-MS, due to evidence of formation of a dimeric species in one of the binding events of this compound in the electron density maps of the crystal structure. Each fragment was diluted 100-fold from the initial stock solution into 50% methanol/50% water. The diluted fragment solution was then directly injected at a rate of 5 μL min^-1^ into the Q-exactive Plus Mass Spectrometer (Thermo Scientific) and measured in negative mode. Raw data were analyzed using X-Calibur Qual browser (Thermo Scientific) to identify the compounds of interest by their accurate mass. The mass spectrometry analysis confirmed that the samples of fragments M-80, M-83, M-91, M-92 and M109 contained the respective sulfonic acids, whereas the corresponding sulfonyl chlorides were not detected and that the M-72 sample contained also a dimeric derivative of fragment M-72 (**Fig. S1**).

### Structure Analysis

The mode of binding of CoA at the CoA-A(HAD/KAT), CoA-B(ECH2) and CoA-C(ECH/HAD) binding sites is provided by the coordinates of PDB ID 7O4R (2.8Å resolution, referred to as the CoA-A structure), 7O4S (2.8Å resolution, referred to as the CoA-B structure) and 7O4T (2.1Å resolution, referred to as the CoA-C structure), respectively (Dalwani *et al*., 2021). The mode of binding of CoA in the ECH and KAT active sites is also provided by these structures and the CoA-C structure (PDB ID 7O4T) is used as the reference coordinate set. The electrostatic potential on the molecular surface of the MtTFE tetramer was calculated and visualized using CCP4MG (McNicholas *et al*., 2011) using the CoA-C structure (PDB ID 7O4T) as the model. In this structure the main chain for all four chains is completely built. To generate the model for the electrostatic surface calculations, first all the ligand and water molecules were deleted, and then disordered missing side chains of arginine, lysine, glutamate and aspartate residues were modelled into the structure. The structures of the MtTFE-fragment complexes were superimposed on the CoA-C structure using the SSM tool (Krissinel & Henrick, 2004) in COOT and the fragment binding with respect to the electrostatic surface was also visualized using CCP4MG (McNicholas *et al*., 2011). **Movie S1** was generated using CCP4MG (McNicholas *et al*., 2011). All structure figures were generated using PyMOL (the PyMOL Molecular Graphics System, Version 2.0 Schrödinger, LLC).

## Results and Discussion

### The 17 structures of the MtTFE fragment complexes identify new binding sites on the surface of MtTFE

Crystallographic fragment screening against a total of 226 fragments was performed for MtTFE using two compound libraries, (a) a library of 96 chemically diverse small molecules from HZB, and (b) a collection of 130 small molecules obtained from the University of Marburg, selected to include a negative charge, as also present in the substrates and reaction intermediates of MtTFE. Of the 226 fragments that were tested, 19 hits, being MtTFE structures with bound fragments, were obtained, 14 of which belonged to the Marburg collection, indicating that the choice of a library that has more negatively charged fragments was more successful for MtTFE. For 2 of the 19 structures the features of the electron density map at the ligand binding sites were insufficient to satisfactorily decide the ligand orientation. For these two structures, the fragment binding sites overlapped with sites that were also identified in structures of other MtTFE-fragment complexes, and therefore these two structures were excluded from further refinement and the structures of 17 fragment binding experiments (5 HZB fragments and 12 Marburg fragments) were selected and used for further analysis (**Table 1, Table S1**). Of these, the structures of 13 fragments were refined to a resolution of 2.8 Å or better, 2 fragments to 2.9 Å and 2 fragments to a low resolution of 3.0 Å and 3.2 Å, respectively. Unlike typical fragment screening campaigns that are carried out to identify the precise binding modes of fragments as starting points for the development of lead candidates for drug discovery purposes, our fragment screening experiments were aimed at identifying low affinity binding sites on the surface of MtTFE. Therefore, data sets of somewhat lower resolutions are also informative and these low resolution structures were also retained for further analysis. Key structure refinement statistics are provided in **Table S2 and Table S3**. All built fragments have a good fit to the electron density map as described in the Methods part (their RSCC values are listed in **Table S4**) and as shown for a representative set in **Table 2**.

**Table 2.**
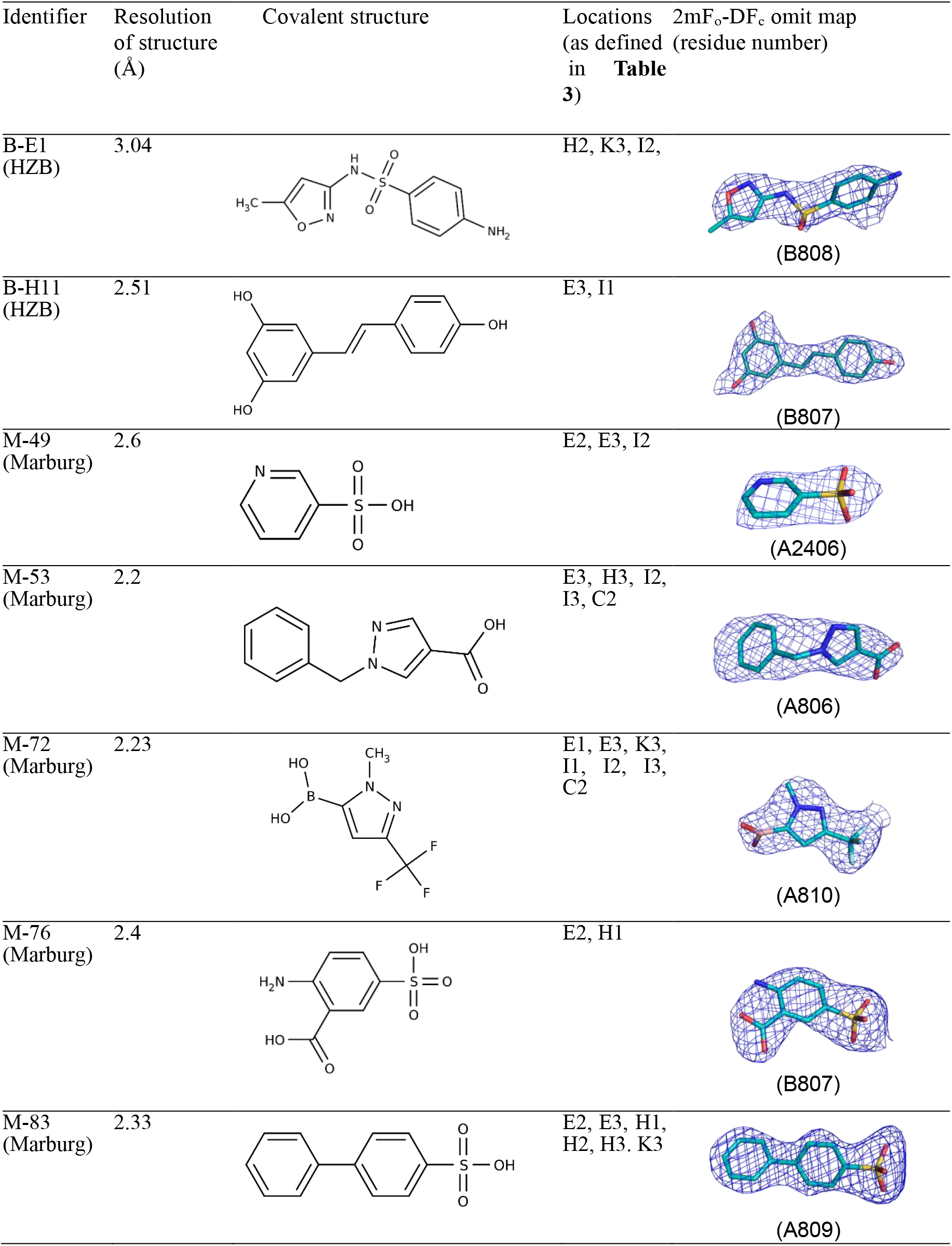
Key information of seven representative fragments that are referred to in the text, being bound at an active site or at the proposed substrate channeling surface near the CoA-C(ECH/HAD) binding site. Further information on these fragments is provided in **Table S1**.

| Identifier | Resolution of structure (Å) | Covalent structure | Locations (as defined in **Table 3**) | 2mF<sub>o</sub>-DF<sub>c</sub> omit map (residue number) |
| --- | --- | --- | --- | --- |
| B-E1 (HZB) | 3.04 |  | H2, K3, I2, | (B808) |
| B-H11 (HZB) | 2.51 |  | E3, I1 | (B807) |
| M-49 (Marburg) | 2.6 |  | E2, E3, I2 | (A2406) |
| M-53 (Marburg) | 2.2 |  | E3, H3, I2, I3, C2 | (A806) |
| M-72 (Marburg) | 2.23 |  | E1, E3, K3, I1, I2, I3, C2 | (A810) |
| M-76 (Marburg) | 2.4 |  | E2, H1 | (B807) |
| M-83 (Marburg) | 2.33 |  | E2, E3, H1, H2, H3, K3 | (A809) |

The 17 fragments are bound over a total of 143 individual binding events. These binding events to a great extent overlap with the previously identified CoA binding sites at the ECH, HAD and KAT active sites as well as with the CoA-A(HAD/KAT), CoA-B(ECH2) and CoA-C(ECH/HAD) binding sites (**Table 3, Table S5)**. Most fragments bind at multiple binding sites (**Table 4**), for example fragment M-1 (25 binding events) and fragment M-76 (20 binding events). In most instances, binding at each site is observed at both copies of the MtTFE tetramer. 118 out of 143 binding events (16 out of 17 fragments) have an aromatic moiety and 129 out of 143 binding events concern negatively charged fragments (12 out of 17 fragments, 9 of which contain a sulfonic acid group) (**Table S1**). Interestingly, although the mother liquor of the crystals used for these experiments contains 2 M (NH_4_)_2_SO_4_ and there are approximately 25 sulfate molecules bound at various surface sites in each structure, the sulfonic acid containing fragments as well as the M-1 fragment (the PF_6_^-1^ anion) do not bind at these sulfate binding sites. Altogether 104 (73%) fragment binding events occur at one of the previously identified binding sites namely the three active sites (including the NAD^+^ binding site), or any of the three additional CoA binding sites or in associated pockets near the bound CoA or NAD^+^ molecules (**Table 4, Table S5**). 39 (27%) individual fragments bind at previously unidentified binding sites, scattered over the surface of the tetramer (**Fig. 2**), 14 of these sites are at crystal contacts. 50 fragment binding events occur at the ECH active site (**Table 4, Table S5**), which is the maximum number of hits at a single binding pocket, indicating that this is the fragment binding “hotspot” of MtTFE. None of the fragments bind in internal cavities and the binding of the fragments has not caused main chain conformational changes of the MtTFE structure.

**Table 3.**
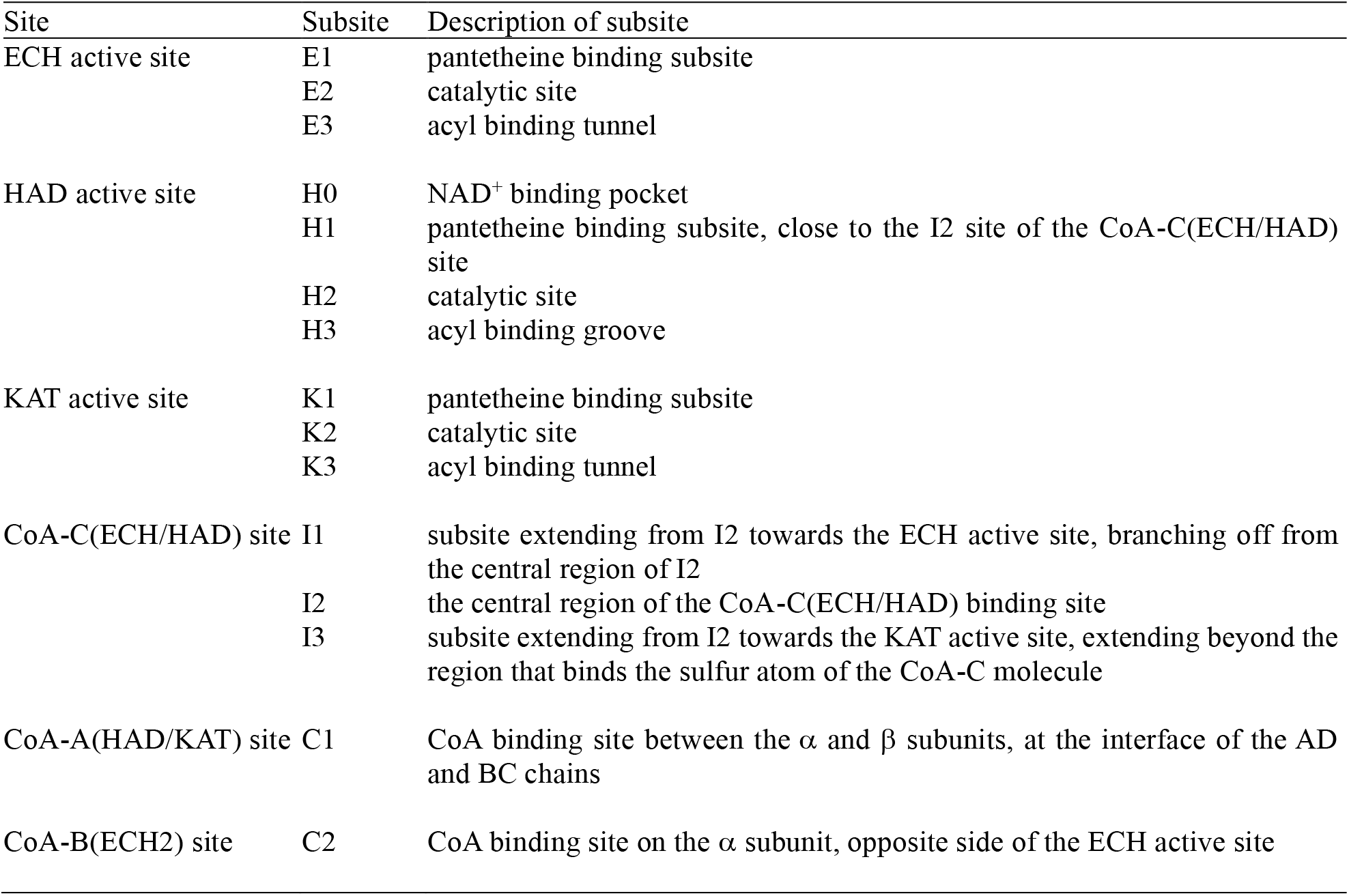
Description of the 6 previously identified CoA binding sites (Dalwani *et al*., 2021) and their subsites.

**Table 4.** Summary of fragment binding events at the 15 fragment binding subsites. The numbers represent the number of binding events associated with the respective binding subsite^a^

|  | ECH active site |  |  | HAD active site |  |  |  | KAT active site |  |  | CoA-C<sup>b</sup> |  |  | CoA-A<sup>b</sup> | CoA-B<sup>b</sup> |  |  |
| --- | --- | --- | --- | --- | --- | --- | --- | --- | --- | --- | --- | --- | --- | --- | --- | --- | --- |
| Subsites | E1 | E2 | E3 | H0 | H1 | H2 | H3 | K1 | K2 | K3 | I1 | I2 | I3 | C1 | C2 | Other | Totals |
| B-E1 |  |  |  |  |  | 2 |  |  |  | 2 |  | 1 |  |  |  |  | 5 |
| B-H11 |  |  | 2 |  |  |  |  |  |  |  | 2 |  |  |  |  |  | 4 |
| B-51 |  |  |  |  |  |  |  |  |  |  |  |  |  | 2 |  |  | 2 |
| B-B3 |  |  |  |  |  |  |  |  |  |  |  |  |  |  |  | 2 | 2 |
| B-77 |  |  |  |  |  |  |  |  |  |  |  |  |  |  |  | 1 | 1 |
| M-1 |  | 2 | 2 | 2 |  | 2 |  |  |  | 2 |  | 6 |  |  |  | 9 | 25 |
| M-10 | 2 |  |  |  |  |  |  |  |  |  |  |  |  |  |  | 1 | 3 |
| M-49 |  | 2 | 1 |  |  |  |  |  |  |  |  | 2 |  |  |  | 3 | 8 |
| M-53 |  |  | 2 |  |  |  | 2 |  |  |  |  | 2 | 2 |  | 1 | 3 | 12 |
| M-72 | 2 |  | 4 |  |  |  |  |  |  | 2 | 1 | 2 | 2 |  | 2 |  | 15 |
| M-76 |  | 2 |  |  | 2 |  |  |  |  |  |  |  |  |  |  | 16 | 20 |
| M-79 |  | 2 |  |  |  |  |  |  |  |  |  |  |  |  |  | 2 | 4 |
| M-80 |  | 2 | 2 | 2 |  |  |  |  |  |  |  |  |  |  |  |  | 6 |
| M-83 |  | 2 | 4 |  | 2 | 2 | 2 |  |  | 2 |  |  |  |  |  | 2 | 16 |
| M-91 |  | 2 | 4 |  |  |  |  |  |  |  |  |  |  |  |  |  | 6 |
| M-92 |  | 2 | 4 |  |  |  |  |  |  |  |  |  |  |  |  |  | 6 |
| M-109 |  | 2 | 3 |  |  | 1 | 2 |  |  |  |  |  |  |  |  |  | 8 |
| Totals | 4 | 18 | 28 | 4 | 4 | 7 | 6 |  |  | 8 | 3 | 13 | 4 | 2 | 3 | 39 | 143 |
| Totals | 50 |  |  | 21 |  |  |  | 8 |  |  | 20 |  |  | 2 | 3 | 39 | 143 |

<sup>a</sup> The subsites are described in **Table 3**. The binding mode of some fragments overlaps with two subsites, in such cases only one subsite is highlighted. In most cases binding events occur in pairs, as they occur in both copies of the asymmetric unit. Detailed information about the fragments is provided in **Table S1** and about the subsites in **Table S5**.
<sup>b</sup> CoA-A, CoA-B and CoA-C refer to the CoA-A(HAD/KAT), CoA-B(ECH2) and CoA-C(ECH/HAD) sites.

**Figure 2.**
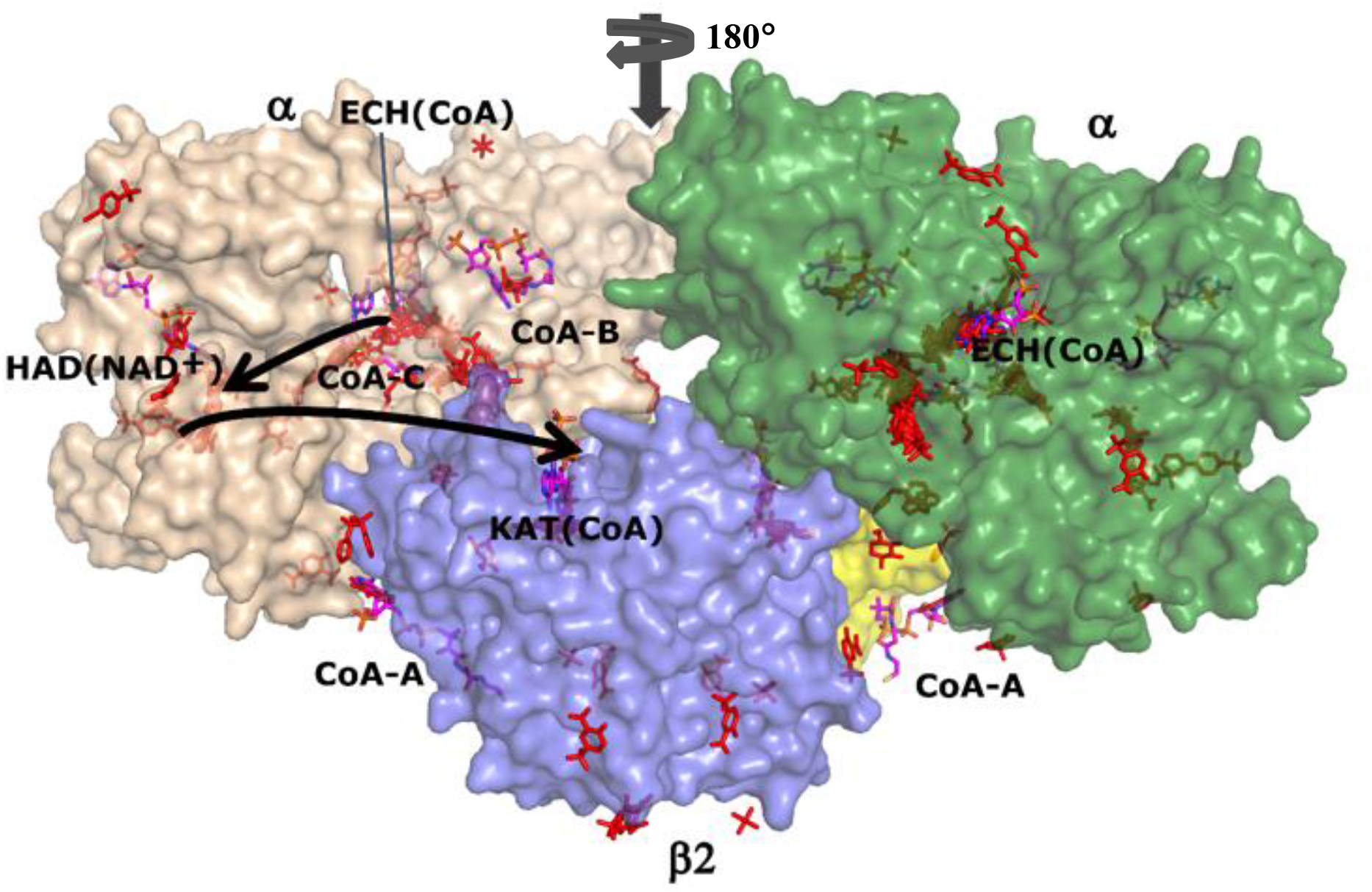
The 143 binding events on the surface of the TFE tetramer. Same view and coloring scheme as used in **Fig. 1**. The two α-subunits are colored wheat and green and the β_2_ thiolase dimer subunits are colored cyan and yellow. The ECH, HAD and KAT active sites are identified as ECH(CoA) (with thin arrow), HAD(NAD^+^) and KAT(CoA). The vertical arrow visualises the twofold axis of the α_2_β_2_ tetramer. The ECH active site of the left α-subunit (wheat) is behind and the ECH active site of the right α-subunit (green) is at the front, showing the fragments bound at its substrate binding tunnel. The surface has been made transparent, such that binding events that are hidden behind the surface are still visible. The labels CoA-A, CoA-B and CoA-C identify the additional CoA binding sites.

Noteworthy, in some structures blobs of positive difference electron density could not be assigned to the used fragments and in these cases the densities have been left un-modelled. This can be for example because of partial disorder of the bound fragment. One fragment (M-72) was observed in its unmodified form (see for example **Table 2**) but the electron density map also showed it to be present in a modified dimeric form having reacted in one case with what appears to be a glycerol and in two other cases with a histidine side chain of the enzyme. The modified histidine side chain is observed in both copies of the N-terminal His-tag of the α-chain, at His (−9), and its binding site is at the exit of the substrate binding tunnel of the ECH active site. The structures of the modelled derivatives of M-72 are in agreement with known structural properties of this boron compound (Diaz & Yudin, 2017). The occurrence of the monomeric and dimeric forms of M-72 in the stock solution was verified by mass spectrometry (see Methods section). The bound unmodified M-72 molecules and the modified M-72 derivatives included in the final structure, respectively, have a good fit with the electron density map, having RSCC values of 0.82 or higher. None of the other compounds resulted in a covalent modification of MtTFE.

### The fragment binding events that are mapped to the 15 subsites, defined by the three active sites and the three additional CoA binding sites

Previous crystallographic binding studies of MtTFE with CoA have identified the ECH, HAD and KAT active sites, as well as three additional CoA binding sites, referred to as the CoA-A(HAD/KAT), CoA-B(ECH2) and CoA-C(ECH/HAD) binding sites, based on their location on the surface of the MtTFE tetramer (Dalwani *et al*., 2021) (**Table 3, Fig. 1**). Each of these six sites occurs in pairs, as the asymmetric unit of this crystal form is the α_2_β_2_ tetramer, in which a local twofold axis (**Fig. 1**) relates the two αβ-subunits to each other. Many binding sites of the fragments overlap with these CoA binding regions, but also identify binding pockets near these CoA binding sites, for example, regions where the acyl-tail of the acyl-CoA substrate molecules could bind, as discussed further below.

The substrates of MtTFE are large molecules (M>900 Da), larger than the bound fragments, and therefore each of the active site CoA binding pockets are described in terms of smaller subsites from the pantetheine region (site-1), via the catalytic site (site-2) to the acyl-tail region (site-3), as defined in **Table 3**: (1) three subsites at the ECH active site: E1, E2 and E3, (2) four subsites at the HAD active site: H0 (the NAD binding site), H1, H2 and H3 and (3) three subsites at the KAT active site: K1, K2, and K3. The CoA-C(ECH/HAD) additional CoA binding site and its nearby regions are also divided in three subsites (I1, I2, and I3, as defined in **Table 3**). Only a few fragments are bound at the other two additional CoA binding sites and therefore these sites are referred to as the C1 and C2 sites for the CoA-A(HAD/KAT) and CoA-B(ECH2) sites, respectively. In this way, 104 fragment binding events are mapped to these15 subsites (**Table 4)**. In addition, 39 other binding events are observed at binding sites scattered over the surface of the MtTFE tetramer (**Fig. 2**). In the next sections the fragment binding in the three active sites is discussed and subsequently the binding events at the three additional CoA binding sites are described.

### Fragment binding at the three active sites include the proposed CoA acyl tail binding pockets

Previous crystallographic binding experiments of seven MtTFE active site point mutated variants with 2*E*-enoyl-CoA substrates did not yield structures of complexes of MtTFE with bound acyl-CoA substrate/intermediate molecules, but instead, resulted in CoA bound structures (Dalwani *et al*., 2021). Thus, although, these previous studies identify the binding sites for the CoA moiety of the acyl-CoA substrate at each of the active sites, as well as binding of CoA at three additional sites, these studies did not provide experimental evidence for the mode of binding of the acyl-tail part of the acyl-CoA substrate in any of the active sites. However, in addition to fragment binding being observed at the CoA binding sites, the current fragment screening experiments resulted also in MtTFE structures where fragments are bound at the predicted acyl tail-binding pockets of each of the active sites – referred to in **Table 3** as E3, H3 and K3. A similar overlap of binding sites for fragments and fatty acids has also been described in recent crystallographic binding studies of serum albumins with ketoprofen (Anderson, 2022; Czub *et al*., 2022) using a fragment of similar properties as used in these MtTFE studies (ketoprofen has a negative charge and two aromatic rings).

### The ECH active site binding pocket

A majority of the fragments (13 out of 17 fragments, 50 binding events) bind in the ECH binding pocket, making it the fragment binding hotspot of our fragment screening experiments (**Table 4**). Multiple copies of each fragment are bound across the entire ECH binding site on both sites of the tetramer (**Fig. 3**). Fragments M-49, M-76, M-79, M-80, M-83, M-91, M-92 and M-109 share a common substructure being a sulfonic acid functional group associated with an aromatic ring. Interestingly, the binding site for the sulfonic acid functional groups (at the E2 site) is the same in all these fragments. Fragments M-91 and M-92 differ only by a single substitution being either a bromide or chloride moiety and 3 molecules of each fragment bind at subsites E2 and E3 such that even the aromatic rings overlap with each other. 2 molecules of fragment M-80 are similarly bound, again overlapping with 2 out of 3 of the M-91 and M-92 fragments. Similarly, fragments M-76 and M-79 differ by only a single substitution at position 2 of the aromatic ring, being either an NH2 or an F functional group, and a common binding site, at subsite E2, is observed. These fragments also superpose on each other. Although their sulfonic acid functional groups occupy the same positions as M-80, M-91 and M-92 (at the E2 site), the benzoic acid moiety of the aromatic rings is flipped, facing the opposite direction. Fragments B-H11 and M-83 bind at the catalytic site (E2) such that they interact directly with the side chains of the catalytic glutamates, αGlu119 and αGlu141. Thus, bound fragment molecules cover the entire ECH active site, from the pantetheine binding site through the substrate binding tunnel to the exit of this tunnel (M72, M83), identifying the substrate binding pocket for the linear tail of 2*E*-enoyl-CoA substrates. The bound fragment molecules nicely overlap with the predicted mode of binding of the tail of 2*E*-decenoyl-CoA from previous modelling experiments (Dalwani *et al*., 2021), as shown in **Fig. 4**, using the bound M-83 fragments as an example. It is noteworthy that none of the fragments bind at the adenine binding pocket, neither at the ECH active site nor at the HAD and KAT active sites.

**Figure 3.**
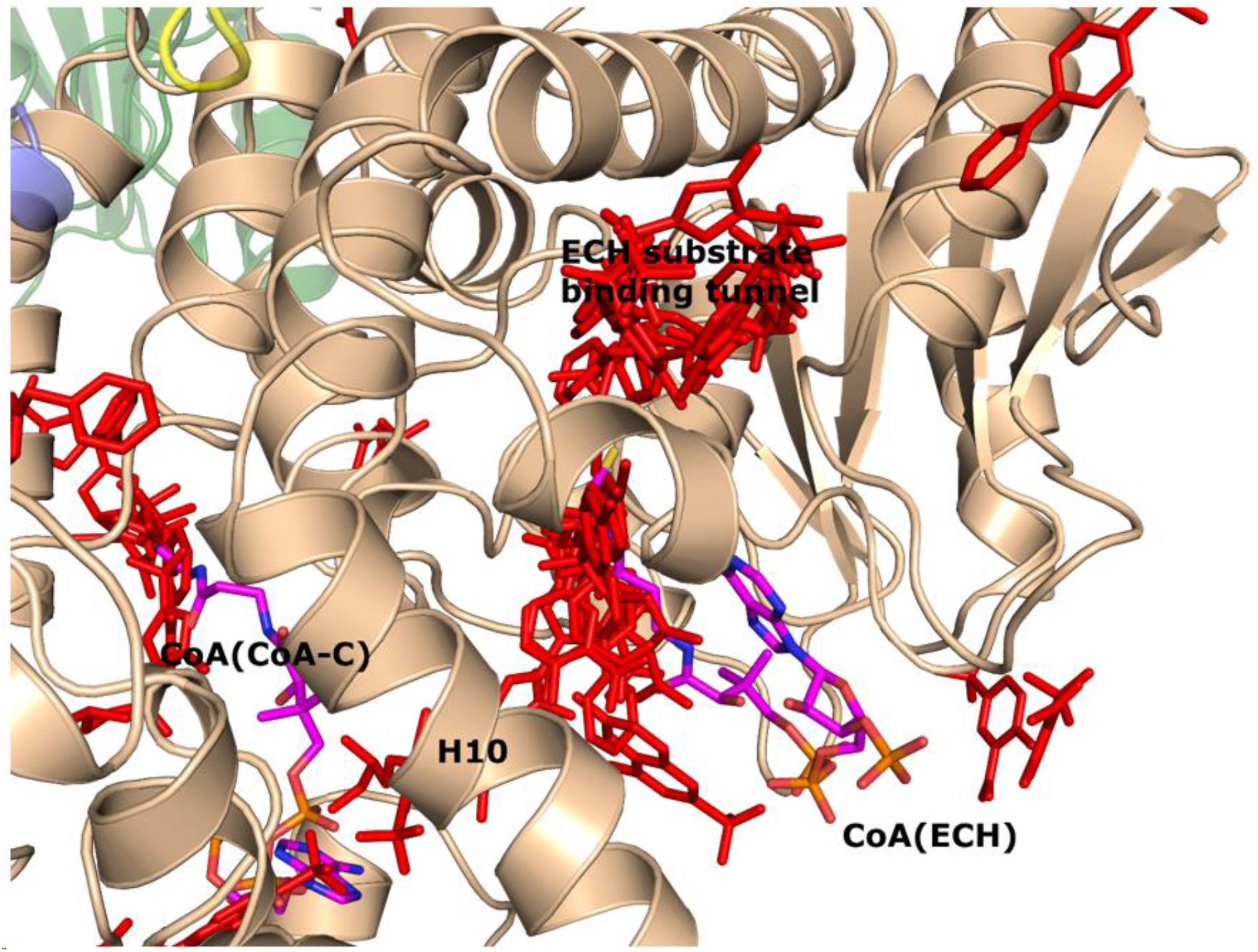
Fragment binding events at the ECH active site. View into the ECH substrate binding tunnel (view is obtained by rotating 180° around the vertical direction of **Fig**. 1). The C-terminal helix of the crotonase fold (helix H10) covers the ECH active site. Also included are CoA as bound to the ECH active site (magenta, labeled as CoA(ECH) as well as the CoA bound at the CoA-C(ECH/HAD) site (magenta, labelled as CoA(CoA-C). There are no fragments bound at the adenine binding site of the CoA(ECH) molecule. The bound fragments cover the protein surface from the CoA-C site to the exit of the ECH substrate binding tunnel.

**Figure 4.**
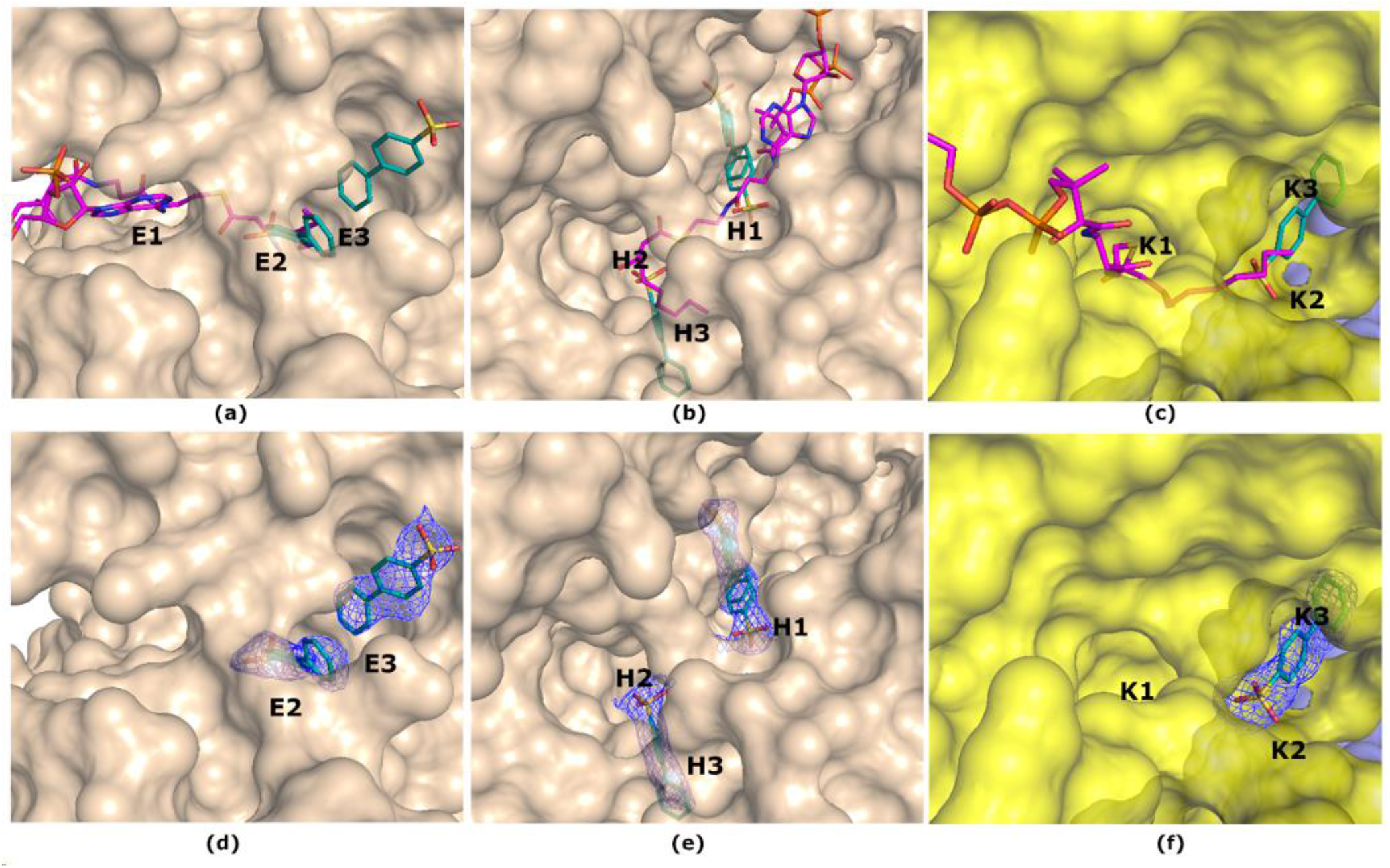
The subsites of the ECH (E1, E2, E3), HAD (H1, H2, H3) and KAT (K1, K2, K3) active sites. (*a, b, c*) The mode of binding of fragment M-83 (cyan) is shown to the E3, H3, and K3 subsites, together with the superimposed mode of binding of the acyl-CoA substrate (magenta), as predicted by model building (Dalwani et al, 2021). (*d, e, f*) The 2mF_o_-DF_c_ omit maps for the M-83 molecules (as visualized in the upper panels) are shown, contoured at 1.0 σ.

### The HAD active site binding pocket

The HAD active site comprises of the C domain (NAD^+^ binding domain) and D/E domains of the α subunit of MtTFE. The C domain is known to exhibit conformational flexibility with respect to the D/E domains such that it can exist in ‘open’ and ‘closed’ conformations. The fully-closed conformation is only captured in the structure of the complex of the homologous monofunctional human HAD, complexed with NAD^+^ and acetoacetyl-CoA, adopting the completely closed conformation, that is the competent form of this enzyme (Barycki *et al*., 2000). In the available structures of MtTFE, the substrate binding groove of the HAD active site is always seen in its open conformation. Except for capturing NAD^+^ binding in the NAD^+^ binding pocket (subsite H0) of the HAD active site (Dalwani *et al*., 2021), all previous crystallographic binding experiments failed to capture any ligand binding in the 3*S*-hydroxyacyl-CoA binding pocket of the HAD active site (i.e. subsites H1, H2 and H3). In our crystallographic fragment screening experiments, 7/17 fragments (21 binding events) bind at the HAD active site (**Table 4**). Fragments B-E1, M-53, M-83 and M-109 form hydrogen bonds either with αSer441 or both αSer441 and αSer512, which both point into the catalytic site. The mode of binding of fragment M-83 is shown in **Fig. 4**. Only the M-1 and M-109 fragments have interactions within 3.7 Å with the side chain of the catalytic αHis462. Even though a lesser number of fragments bind here as compared to the ECH active site, similar to the ECH active site, binding is seen at all subsites of this active site (**Table 4**), such that the bound fragment molecules overlap also with the predicted mode of binding of 3-ketodecanoyl-CoA at the HAD active site of MtTFE (Dalwani *et al*., 2021). None of the bound fragments induce conformational changes that capture a closed conformation of the HAD active site in MtTFE. Thus, in all the structures of MtTFE reported this far, the HAD active site of MtTFE is always seen in an open conformation.

### The KAT active site binding pocket

The acyl-tail binding pocket of the KAT active site is the least accessible of the three active sites of MtTFE. Least number of fragments bind at the KAT active site (4/17 fragments are bound, 8 binding events) (**Table 4**). Each of these bound fragments (B-E1, M-1, M-72 and M-83) bind in the KAT acyl-tail binding tunnel, beyond the CoA sulfhydryl group and they do not interact with the side chain of the catalytic cysteine, βCys92. Interestingly, the R-SO_2_-R and R-SO_2_-OH functional groups of B-E1 and M-83, respectively bind at the same site, forming hydrogen bonds with the side chain of βGln149. M-1 (the hexafluorophosphate anion), and the trifluoro functional group of M-72 bind also at this site, in a different mode, displacing the side chain of βMet134, which has moved. The mode of binding of fragment M-83 is shown in **Fig. 4**. The bound fragments overlap with the predicted mode of binding of the tail of 3-ketodecanoyl-CoA from previous modelling experiments (Dalwani *et al*., 2021).

### The fragment binding events identify CoA-C(ECH/HAD) as a possible functional transient binding site

The reaction intermediates of MtTFE are polar negatively charged derivatives of CoA. The analysis of the surface features of MtTFE shows that its three active sites are separated by a surface path which is devoid of negatively charged residues (**Fig. 5, Fig. S2, Movie S1**) and it has been proposed that this path could be used to transiently anchor the reaction intermediates allowing them to crawl between the active sites, thus enabling substrate channeling (Venkatesan & Wierenga, 2013). Crystallographic binding studies with CoA have identified three additional CoA binding sites (Dalwani *et al*., 2021), which to some extent overlap with this surface path. A comparison of the CoA-protein interactions of CoA bound at these additional binding sites versus the CoA-protein interactions of CoA bound in the active sites shows that on average, the total number of atom-atom interactions per bound CoA (23.3 versus 27.5) and the total number of hydrogen bond interactions (3.2 versus 8.5) are significantly less for CoA bound in the additional binding sites, as compared to CoA bound in the active site binding pockets (**Table S6**). This is in particular interesting for the CoA-C(ECH/HAD) binding site, which is located between the ECH, HAD and KAT active sites and which has been proposed to be part of the surface path relevant for substrate channeling (Dalwani *et al*., 2021). At this binding site the CoA is bound in an extended conformation and the interactions between the bound CoA and the protein are much weaker than at the active sites (**Table S6**). The location of this site and the weak CoA-protein interactions indicate that of the three additional binding sites, the CoA-C(ECH/HAD) site could be a functional transient binding site for reaction intermediates.

**Figure 5.**
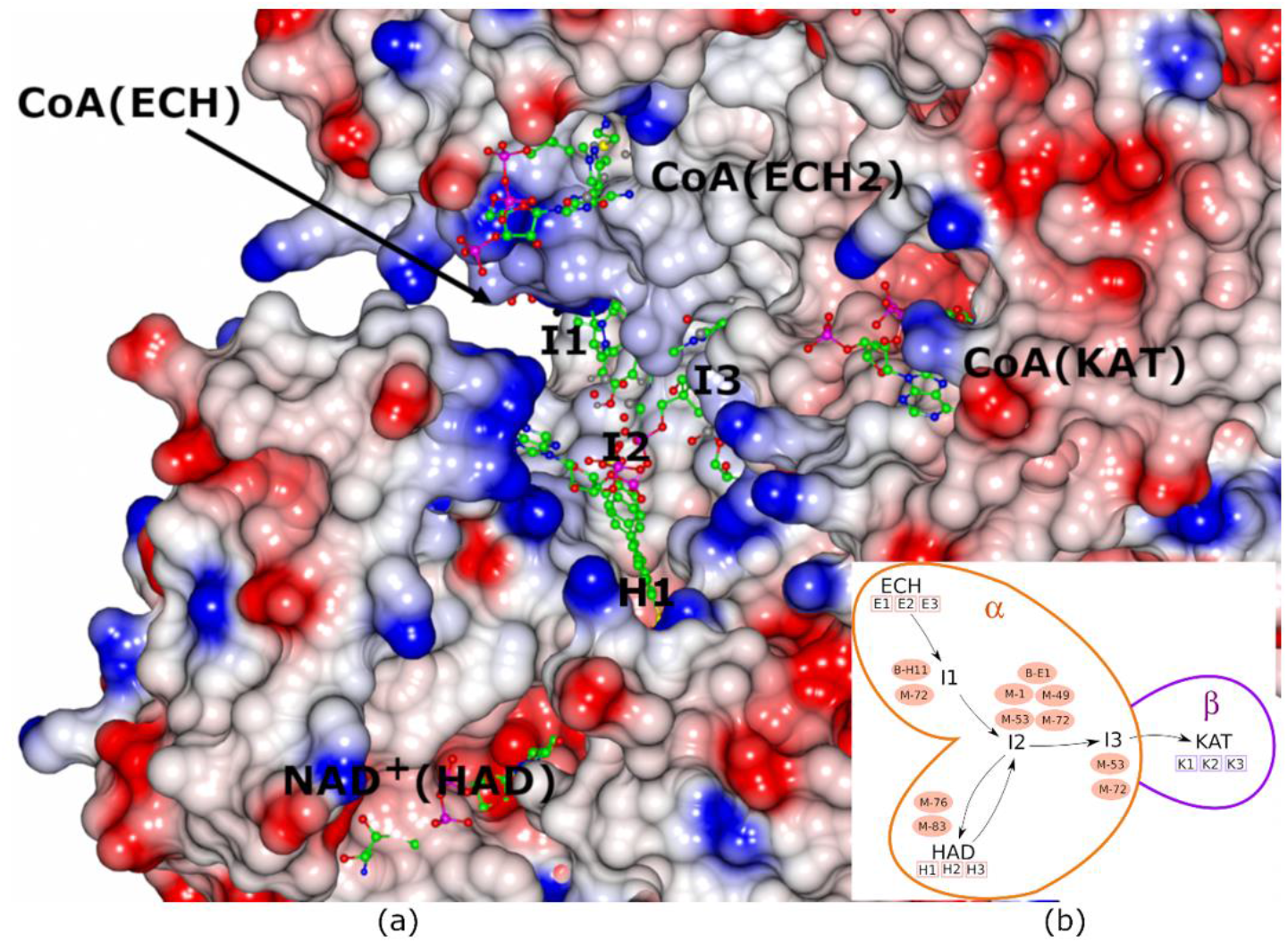
Properties of the protein surface of the possible substrate channeling path between the three active sites of MtTFE. The image shows the electrostatic molecular surface of the CoA-C(ECH/HAD) region, near the interface of chain A (α subunit) and chain D (β subunit) of the CoA-C structure (PDB ID 7O4T), color coded such that the blue and red colors identify surface regions with positive and negative electrostatic potential, respectively. Neutral regions have a white color. The active sites are labelled as CoA(ECH) (with arrow), NAD^+^(HAD) (of the α subunit) and CoA(KAT) (of the β subunit). The CoA molecule bound at the CoA-C(ECH/HAD) site (I2) at the center of the surface between the three active sites is also included. I1, H1 and I3 identify the binding regions extending from I2 towards the ECH, HAD and KAT active sites, respectively. The fragments shown are indicated in the schematic view (insert). A stereo figure is shown in **Fig. S2** and a video (**movie S1**) is also provided as Supplementary Material.

The fragment screening experiments identify multiple binding sites which overlap with the additional CoA binding sites. 6/17 fragments (18 binding events) bind overlapping with at least one of the three additional CoA binding sites (**Table 4**). A single fragment (B-51) binds at CoA-A(HAD/KAT) (2 binding events), 2 fragments (M-53 and M-72) bind at CoA-B(ECH2) (3 binding events) and 5 fragments (B-E1, M-1, M-49, M-53, and M-72) bind at the CoA-C(ECH/HAD) binding site (I2: 13 binding events). Clearly, the CoA-C(ECH/HAD) is a favorable binding site for these fragments (**Fig. 5**). Furthermore, near the CoA-C(ECH/HAD) site, fragments B-H11 and M-72 bind at subsite I1, which perfectly bridges the gap between the bound CoA at the ECH active site and the CoA-C(ECH/HAD) binding site. In addition, fragment M-76 and M-83 bind at H1 (the binding region of the pantetheine moiety of the HAD active site, which is close to I2) and fragments M-53 and M-72 bind at subsite I3, which is extending beyond the CoA sulfur atom, towards the KAT active site. **Fig. 5** provides also a schematic visualisation of the I1, H1 and I3 binding sites near the CoA-C(ECH/HAD) site (I2) with respect to the ECH, HAD and KAT active sites. The negatively charged functional groups of fragments M-1 and M-83, which bind at subsites H1 and I2, overlap with the negatively charged pyrophosphate moiety of the CoA molecule bound at this site. Each of the fragments that bind at the CoA-C(ECH/HAD) binding region also bind to at least one of the three active sites (**Table 4**). More specifically, fragments M-53 and M-72 which bind at the I2 and I3 subsites also bind at the E3 and H3 subsites (M-53) as well as at the E3 and K3 subsites (M-72).

### The proposed substrate channeling path identified by the fragment binding events

As outlined above, in addition to identifying the CoA-C(ECH/HAD) site as a potential transient binding site (the I2 site) for reaction intermediates, our crystallographic fragment screening experiments identify subsites I1, H1 and I3 which are binding pockets that extend from the CoA-C binding site (I2) to the ECH, HAD and KAT catalytic sites, respectively. The crystallographic fragment screening experiments suggest therefore a substrate channeling path that can be used for the surface crawling of negatively charged reaction intermediates between the three active sites. The proposed path consists of the binding sites ECH (catalytic site) → I1 → I2 → H1 → HAD (catalytic site) → H1 → I2 → I3 → KAT (catalytic site). The distances from the ECH and HAD active sites to the I2 site are approximately 21Å and 12Å, respectively and from the KAT active site the corresponding distance is approximately 44Å, when considering the distances between the bridging pyrophosphate oxygen atoms of the CoAs bound at these sites. The path is identified by the fragments B-E1, B-H11, M-1, M-49, M-53, M-72, M-76 and M-83, in addition to the CoA-C(ECH/HAD) molecule (**Fig. 5, Fig. S2**). The **Movie S1** visualises the results of these fragment binding experiments, highlighting the proposed substrate channeling path between the active sites, by showing the fragments as bound at sites along this surface path of MtTFE. The fragments, M-1, M-49, M-53, M-72, M-76 and M-83 are negatively charged, like the MtTFE reaction intermediates. This path can be described as consisting of neutral residues, being lined by the side chains of positively charged residues (Lys/Arg) (**Fig. 5**). Positively charged residues of the protein surface have been proposed to be important for steering the diffusion of negatively charged ligands for substrate channeling of the TS-DHFR bifunctional enzyme (Anderson, 2017; Metzger *et al*., 2014) as well as in other biological systems (Zheng *et al*., 2019). Other properties, such as disfavouring the binding of water molecules, referred to as “dewetting” (Hilario *et al*., 2016) have been described for the substrate channeling path of the enzyme tryptophan synthase in which a neutral indo le molecule channels through a tunnel between its two active sites.

### Concluding remarks

The crystal form of unliganded MtTFE used in these studies has several advantages to identify new, weak affinity binding sites on the surface of MtTFE by the fragment binding approach, using crystal soaking. In this crystal form the asymmetric unit is the MtTFE tetramer, which means that there are two copies of each unique binding region per asymmetric unit. The percentage solvent of this crystal form is relatively high (V_M_ is 3.9 Å^3^/Da, which corresponds to 69% solvent) meaning that the molecules are loosely packed and therefore large portions of the surface of the protein are not involved in crystal packing interactions, but instead can interact with solutes. Indeed, most binding events occur as pairs, being observed in both copies present in the asymmetric unit (**Table 4**). These fragment binding studies show preferred binding events in the active sites, including in the predicted acyl tail binding pockets (**Fig. 4**), as well as in regions between the active sites (**Fig. 5**). 16 (out of 17) of the characterized MtTFE-fragment complexes concern compounds with aromatic rings (**Table S1**) (the only exception is fragment M1, which is the PF_6_^-1^ anion). 12 (out of 17) fragments are negatively charged anions in solution. 79 fragment binding events occur in pockets that shape the three active sites (**Table 4**), of which 42 binding events concern the binding pockets for the acyl tails. A very striking observation concerns the 24 binding events in the region of the CoA-C binding site (I2), extending to the ECH, HAD and KAT active sites (I1, H1, I3) (**Fig. 5, Fig. S2, Movie S1**). The visualisation of the electrostatic surface properties indicates that this region is devoid of negative charges. The presence of a transient binding site (I2), extended by the binding sites I1, H1, and I3 to the ECH, HAD and KAT active sites, respectively, is consistent with a substrate channeling mechanism of reaction intermediates between these active sites by surface crawling, just as described also in the binding studies of the *Mtb* tryptophan synthase (Bosken *et al*., 2022; D’Amico & Boehr, 2023) and the TS-DHFR bifunctional enzymes (Anderson, 2017; Metzger *et al*., 2014). Thus, this MtTFE study provides a basis for further studies to better characterize the substrate channeling properties of MtTFE. Orthogonal approaches would be required for such studies, in addition to these crystallographic binding experiments, to discover for which acyl-CoA substrates channeling is relevant in MtTFE and also to validate the proposed path of channeling.

## Acknowledgements

The data sets for these studies were collected at PETRA III (beam line P13), DLS (beam lines I03, I04-1), MAX IV (beam line BioMAX), and BESSY II (beam line 14.1) and we thank the beam line scientists of these beam lines for their support. We gratefully acknowledge the help of Dr. Andrey Lebedev (CCP4) with making the restraint file of the modified histidine side chain, as observed in the M-72 fragment structure. The use of the facilities and expertise of the Biocenter Oulu Structural Biology core facility, a member of Biocenter Finland, Instruct-ERIC Centre Finland and FINStruct, is gratefully acknowledged. We thank Gerhard Klebe, Martin Schlitzer, Thorsten Steinmetzer, Sandrine Marchais-Oberwinkler and Ahmed Merabet from the Institute of Pharmaceutical Chemistry at Philipps-University Marburg for providing compound samples. We also acknowledge the support of the mass spectrometry and biophysical characterization core facilities of Biocenter Oulu. This research has been supported by the Academy of Finland (grants 289024, 293369, 319194) and the Magnus Ehrnrooth Foundation.

