## Supplementary material for "Crystallographic fragment binding studies of the *Mycobacterium tuberculosis* trifunctional enzyme suggest binding pockets for the tails of the acyl-CoA substrates at its active sites and a potential substrate channeling path between them": Supplemenetary data

**Supplementary Table S1.** Summary of the molecular properties of the 17 fragments for which binding has been observed. The 5 HZB compounds (B-XX) are hits from the first screen and the 12 Marburg compounds (M-XX) have been obtained from the second screen.

| Identifier | Resolution of structure | Mw(Da) | Predicted overall charge of predominant form at pH 7 | Structural formulas | Number of molecules bound | Stock solution |
|---|---|---|---|---|---|---|
| B-E1 (HZB) | 3.04 | 253.28 | 0 | 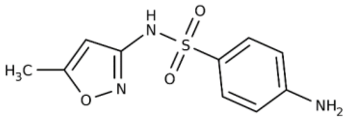 | 5 | 1M in 100% DMSO (pre-spotted)[a] |
| B-H11 (HZB) | 2.51 | 194.19 | 0 | 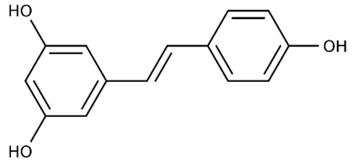 | 4 | 1M in 100% DMSO (pre-spotted)[a] |
| B-51 (HZB) | 2.8 | 228.24 | 0 | 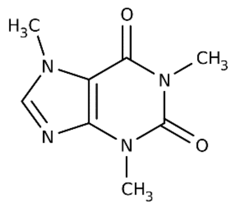 | 2 | 1M in 100% DMSO (pre-spotted)[a] |
| B-B3 (HZB) | 2.9 | 342.3 | 0 | 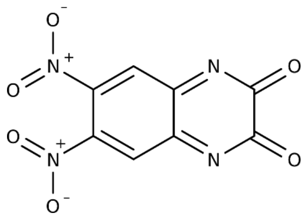 | 2 | 1M in 100% DMSO (pre-spotted)[a] |
| B-77 (HZB) | 2.45 | 252.14 | 0 | 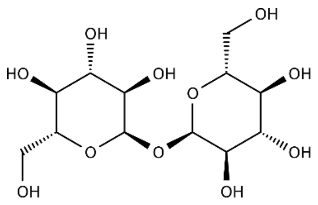 | 1 | 1M in 100% DMSO (pre-spotted)[a] |
| M-1 (Marburg) | 2.7 | 144.96 | -1 | 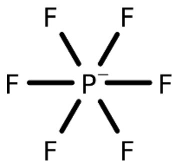 | 25 | 0.5M in 100% DMSO |
| M-10 (Marburg) | 3.2 | 239.24 | -1 | 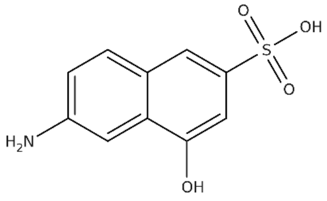 | 3 | 0.5M in 50% DMSO |

| | | | | | | |
|---|---|---|---|---|---|---|
| M-49 (Marburg) | 2.6 | 159.17 | -1 | 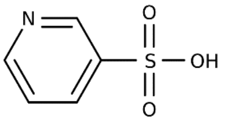 | 8 | 0.5M in 50% DMSO |
| M-53 (Marburg) | 2.2 | 202.21 | -1 | 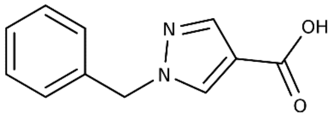 | 12 | 0.5M in 100% DMSO |
| M-72 (Marburg) | 2.23 | 193.92 | -1 | 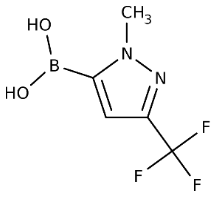 | 15 | 0.5M in 100% DMSO |
| M-76 (Marburg) | 2.19 | 217.2 | -2 | 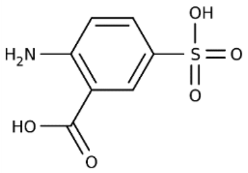 | 20 | 0.5M in 100% DMSO |
| M-79 (Marburg) | 2.59 | 238.62 | -2 | 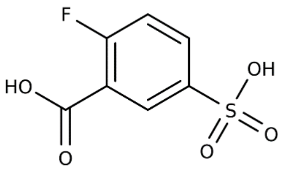 | 4 | 0.5M in 100% DMSO |
| M-80 (Marburg) | 2.4 | 183.19 | -1 | 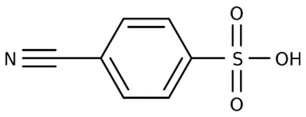 | 6 | 0.5M in 100% DMSO |
| M-83 (Marburg) | 2.33 | 234.27 | -1 | 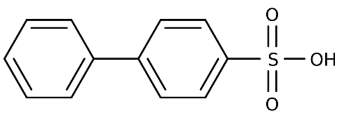 | 16 | 0.25M in 50% DMSO |
| M-91 (Marburg) | 2.62 | 237.07 | -1 | 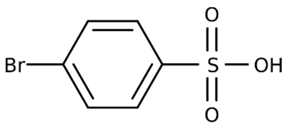 | 6 | 0.5M in aqueous solution |
| M-92 (Marburg) | 2.89 | 192.62 | -1 | 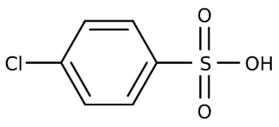 | 6 | 0.5M in aqueous solution |

| | | | | | | |
|---|---|---|---|---|---|---|
| M-109 (Marburg) | 2.77 | 203.17 | -1 | 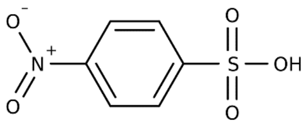 | 8 | 0.5M in aqueous solution |

[a] For these experiments pre-spotted plates were used, as provided by HZB.

**Supplementary Table S2.** Data collection and refinement statistics for the HZB fragments.

| Data set | B-E1 | B-H11 | B-51 | B-B3 | B-77 |
|---|---|---|---|---|---|
| **Data collection** | | | | | |
| Beam line | BESSY II (14.1) | BESSY II (14.1) | BESSY II (14.1) | BESSY II (14.1) | BESSY II (14.1) |
| Detector | Pilatus 6M | Pilatus 6M | Pilatus 6M | Pilatus 6M | Pilatus 6M |
| Wavelength (Å) | 0.8950 | 0.8950 | 0.8950 | 0.8950 | 0.8950 |
| Temperature (K) | 100 | 100 | 100 | 100 | 100 |
| **Data Processing** | | | | | |
| Space group | C2 | C2 | C2 | C2 | C2 |
| Unit cell parameters | | | | | |
| a,b,c (Å) | 248.76, 135.08, 119.74 | 249.69, 134.68, 119.48 | 249.51, 134.50, 118.98 | 248.41, 135.16, 119.16 | 249.83, 134.53, 118.84 |
| α,β,γ ($^{O}$) | 90.0, 110.61, 90.0 | 90.0, 110.56, 90.0 | 90.0, 110.86, 90.0 | 90.0, 110.56, 90.0 | 90.0, 110.82, 90.0 |
| Processing software | xdsapp [1], [2] | xdsapp [1], [2] | xdsapp [1], [2] | xdsapp [1], [2] | xdsapp [1], [2] |
| Resolution range (Å) | 48.42-3.04 (3.23-3.04) | 48.38-2.8 (2.97-2.80) | 48.19-2.51 (2.67-2.51) | 48.37-2.9 (3.07-2.90) | 48.21-2.45 (2.59-2.45) |
| $R_{pim}$ (%) (all I+/I-) | 14.4 (77.9) | 10.5 (87.6) | 12.7 (172.6) | 7.9 (60.2) | 8.9 (171.1) |
| $CC_{1/2}$ (%) | 98.4 (53.8) | 98.8 (50.4) | 99.0 (14.4) | 99.4(63.9) | 99.5 (20.0) |
| <I/σ(I)> | 6.48 (1.32) | 6.88 (1.14) | 6.00 (0.49) | 10.24 (1.54) | 7.42 (0.53) |
| Completeness (%) | 99.4 (98.1) | 99.1 (98.2) | 97.5 (89.6) | 97.7 (97.1) | 98.2 (94.4) |
| Multiplicity | 6.4 (6.4) | 4.8 (5.0) | 3.8 (3.4) | 3.9 (3.9) | 3.8 (3.5) |
| No. of reflections | 450495 | 439437 | 462833 | 309390 | 505899 |
| No. of unique reflections | 70733 | 90705 | 121668 | 80269 | 133151 |
| Wilson *B* factor ($Å^2$)$^{\#}$ | 65.4 | 61.5 | 57.2 | 59.4 | 57.6 |
| **Refinement Statistics** | | | | | |
| Resolution range (Å) | 46.59 - 3.05 | 48.38 - 2.8 | 48.19 - 2.52 | 48.37 - 2.9 | 47.0 - 2.45 |
| No. of used reflections | 70654 | 90660 | 120658 | 80267 | 132306 |
| $R_{work}$ (%) | 20.97 | 19.80 | 20.84 | 19.85 | 21.08 |
| $R_{free}$ (%) | 24.58 | 23.31 | 23.98 | 22.90 | 24.53 |
| Total No. of atoms | 17099 | 17085 | 17104 | 16868 | 17085 |
| No. of waters | 89 | 155 | 182 | 47 | 140 |
| Average B factors ($Å^2$) | | | | | |
| Protein | 70.9 | 68.1 | 66.6 | 69.8 | 65.2 |
| Others | 95.8 | 97.2 | 101.2 | 104.7 | 101.6 |
| Waters | 55.4 | 53.4 | 52.7 | 51.6 | 52.7 |
| R.m.s. deviations | | | | | |
| Bond lengths (Å) | 0.0015 | 0.0019 | 0.0018 | 0.0021 | 0.0020 |
| Bond angles ($^{O}$) | 0.42 | 0.49 | 0.46 | 0.49 | 0.49 |
| Ramachandran plot (%)[3] | | | | | |

| | | | | | |
|---|---|---|---|---|---|
| Favored | 95.80 | 95.67 | 96.74 | 96.46 | 97.01 |
| Allowed | 4.06 | 4.02 | 3.09 | 3.45 | 2.81 |
| Outliers | 0.13 | 0.31 | 0.18 | 0.09 | 0.18 |
| **PDB ID** | 8OPU | 8OPV | 8OPW | 8OPX | 8OPY |

[#] As reported in the PDB validation report

**Supplementary Table S3.** Data collection and refinement statistics for the Marburg fragments.

| Data set | M-1 | M-10 | M-49 | M-53 | M-72 | M-76 |
|---|---|---|---|---|---|---|
| **Data collection** | | | | | | |
| Beam line | MAX IV (BioMAX) | MAX IV (BioMAX) | Petra (P13) | Petra (P13) | DLS (I04-1) | DLS (I03) |
| Detector | Eiger 16M Hybrid-pixel | Eiger 16M Hybrid-pixel | PILATUS 6M | PILATUS 6M | Pilatus 6M-F | Eiger2 XE 16M |
| Wavelength (Å) | 0.968619 | 0.968619 | 0.9198 | 0.9198 | 0.91589 | 0.976254 |
| Temperature (K) | 100 | 100 | 100 | 100 | 100 | 100 |
| **Data Processing** | | | | | | |
| Space group | C2 | C2 | C2 | C2 | C2 | C2 |
| Unit cell parameters | | | | | | |
| a,b,c (Å) | 249.46, 134.25, 118.4 | 247.92, 134.91, 119.21 | 250.78, 136.14, 121.05 | 247.31, 134.03, 116.41 | 250.96, 134.61, 119.97 | 250.93, 132.8, 119.29 |
| α,β,γ ($^{O}$) | 90.0, 110.74, 90.0 | 90.0, 110.62, 90.00 | 90.0, 110.42, 90.0 | 90.0, 109.69, 90.0 | 90.0, 110.44, 90.0 | 90.0, 110.46, 90.0 |
| Processing software | xds [4], AIMLESS [5] | xds [4], AIMLESS [5] | xds [4], AIMLESS [5] | xds [4], AIMLESS [5] | autoPROC[6], STARANISO[7] | autoPROC[6], STARANISO[7] |
| Resolution range (Å) | 48.32-2.7 (2.75-2.7) | 48.29-3.20 (3.28-3.2) | 117.8-2.6 (2.64-2.6) | 116.42-2.2 (2.24-2.2) | 112.42-2.23 (2.47-2.24) | 117.55-2.19 (2.35-2.19) |
| $R_{pim}$ (%) (all I+/I-) | 4.6 (59.1) | 12 (125.3) | 4.8 (65.6) | 4.7 (85.1) | 4.4 (46.3) | 4.1 (53.5) |
| $CC_{1/2}$ (%) | 99.8 (53.8) | 98.6 (24.1) | 99.6 (45.3) | 99.8 (34.5) | 99.8 (62.3) | 99.9 (55) |
| $\langle I/\sigma(I) \rangle$ | 10.9 (1.3) | 5.9 (1.1) | 9.3 (1.0) | 9 (1.1) | 11.5 (1.6) | 13.3 (1.5) |
| Completeness (%) | 99.7 (95.9) | 99.5 (97) | 99.6 (92.1) | 99.7 (94.9) | 95.7 (74) | 95.3 (65.3) |
| Multiplicity | 4.8 (4.8) | 3.8 (3.8) | 6.7 (6.2) | 6.8 (6.6) | 5.3 (5.6) | 7.0 (7.0) |
| No. of reflections | 486278 | 228561 | 784947 | 1223214 | 701657 | 1017298 |
| No. of unique reflections | 100954 | 60500 | 116826 | 179810 | 131177 | 145082 |
| Wilson *B* factor (Å$^2$)[#] | 71.6 | 94.9 | 71.1 | 52.4 | 46.8 | 46.5 |
| **Refinement Statistics** | | | | | 46.8 | |
| Resolution range (Å) | 48.32 - 2.7 | 48.29 - 3.2 | 58.75 - 2.6 | 48.98 - 2.2 | 112.42 - 2.24 | 27.08 - 2.19 |
| No. of used reflections | 100936 | 60087 | 116687 | 179610 | 131169 | 145008 |
| $R_{work}$ (%) | 18.24 | 20.39 | 19.16 | 19.30 | 18.62 | 19.14 |
| $R_{free}$ (%) | 21.83 | 24.25 | 22.71 | 22.16 | 22.05 | 21.94 |
| Total No. of atoms | 17164 | 16883 | 16676 | 17257 | 17660 | 17691 |
| No. of waters | 77 | 68 | 196 | 190 | 502 | 596 |
| Average B factors (Å$^2$) | | | | | | |
| Protein | 73.3 | 105.4 | 82.2 | 69.7 | 57.9 | 56.8 |

| | | | | | | |
|---|---|---|---|---|---|---|
| Others | 110.8 | 129.9 | 116.7 | 96.6 | 83.0 | 86.6 |
| Waters | 59.8 | 72.1 | 66.6 | 52.0 | 48.9 | 51.1 |
| R.m.s. deviations | | | | | | |
| Bond lengths (Å) | 0.0020 | 0.0020 | 0.0017 | 0.0019 | 0.0022 | 0.0025 |
| Bond angles ($^O$) | 0.47 | 0.46 | 0.46 | 0.51 | 0.54 | 0.55 |
| Ramachandran plot (%) | | | | | | |
| Favored | 96.77 | 96.73 | 96.50 | 96.30 | 96.93 | 96.70 |
| Allowed | 3.14 | 3.13 | 3.23 | 3.53 | 2.98 | 3.12 |
| Outliers | 0.09 | 0.13 | 0.27 | 0.18 | 0.09 | 0.18 |
| **PDB ID** | 8OQL | 8OQM | 8OQO | 8OQN | 8PF8 | 8OQP |

[#] As reported in the PDB validation report

**Supplementary Table S3.** Data collection and refinement statistics for the Marburg fragments (continued).

| Data set | M-79 | M-80 | M-83 | M-91 | M-92 | M-109 |
|---|---|---|---|---|---|---|
| **Data collection** | | | | | | |
| Beam line | MAX IV (BioMAX) | MAX IV (BioMAX) | MAX IV (BioMAX) | MAX IV (BioMAX) | MAX IV (BioMAX) | MAX IV (BioMAX) |
| Detector | Eiger 16M Hybrid-pixel | Eiger 16M Hybrid-pixel | Eiger 16M Hybrid-pixel | Eiger 16M Hybrid-pixel | Eiger 16M Hybrid-pixel | Eiger 16M Hybrid-pixel |
| Wavelength (Å) | 0.976254 | 0.976254 | 0.976254 | 0.976254 | 0.976254 | 0.976254 |
| Temperature (K) | 100 | 100 | 100 | 100 | 100 | 100 |
| **Data Processing** | | | | | | |
| Space group | C2 | C2 | C2 | C2 | C2 | C2 |
| Unit cell parameters | | | | | | |
| a,b,c (Å) | 250.34, 134.02, 119.75 | 250.70, 135.61, 120.89 | 250.46, 134.70, 119.30 | 247.48, 133.23, 115.93 | 249.46, 134.89, 119.48 | 248.79, 135.63, 119.69 |
| α,β,γ ($^O$) | 90.0, 110.53, 90.0 | 90.0, 110.27, 90.0 | 90.0, 110.68, 90.0 | 90.0, 109.65, 90.0 | 90.0, 110.47, 90.0 | 90.0, 110.63, 90.0 |
| Processing software | autoPROC[6], STARANISO[7] | autoPROC[6], STARANISO[7] | autoPROC[6], STARANISO[7] | autoPROC[6], STARANISO[7] | autoPROC[6], STARANISO[7] | edna_proc, STARANISO[7] |
| Resolution range (Å) | 117.2-2.59 (2.82-2.59) | 117.59-2.4 (2.64-2.4) | 117.16-2.33 (2.49-2.33) | 116.54-2.62 (3.01-2.62) | 116.86-2.89 (3.17-2.89) | 45.51-2.77 (2.9-2.77) |
| $R_{pim}$ (%) (all I+/I-) | 11.9 (143.2) | 11.8 (126.6) | 6.1 (49.3) | 13.2 (77.6) | 7.9 (57.1) | 7.8 (74.9) |
| $CC_{1/2}$ | 98 (25.1) | 98.2 (33.9) | 99.4 (59.1) | 97.7 (30.1) | 99.2 (49.8) | 99.3 (41.2) |
| $\langle I/\sigma(I) \rangle$ | 7.2 (1.5) | 5.9 (1.5) | 9.1 (1.5) | 6.1 (1.7) | 8.5 (1.5) | 8.3 (1.1) |
| Completeness (%) | 92.2 (53.7) | 91.8 (52.1) | 92.9 (60.9) | 91.7 (67.5) | 93.2 (65.7) | 91.9 (52.3) |
| Multiplicity | 6.7 (7.2) | 6.5 (7.2) | 7.1 (6.2) | 7.0 (7.2) | 6.9 (7.3) | 6.7 (6.8) |
| No. of reflections | 601604 | 773421 | 801947 | 409060 | 446439 | 540036 |
| No. of unique reflections | 89654 | 118277 | 113738 | 58657 | 64333 | 80655 |
| Wilson *B* factor ($Å^2$)[#] | 55.1 | 48 | 40.2 | 44.3 | 77.7 | 70.5 |
| **Refinement Statistics** | | | | | | |
| Resolution range (Å) | 50.27 - 2.59 | 47.92 - 2.4 | 55.81 - 2.33 | 54.59 - 2.62 | 116.86 - 2.89 | 45.51 - 2.78 |

| | | | | | | |
|---|---|---|---|---|---|---|
| No. of used reflections | 89611 | 118191 | 113689 | 58621 | 64297 | 80594 |
| $R_{work}$ (%) | 19.10 | 21.82 | 17.83 | 19.89 | 17.97 | 19.95 |
| $R_{free}$ (%) | 23.0 | 26.03 | 21.63 | 24.55 | 21.72 | 23.38 |
| Total No. of atoms | 17000 | 17221 | 18026 | 17002 | 16971 | 16917 |
| No. of waters | 192 | 454 | 719 | 81 | 71 | 50 |
| Average B factors (Å$^2$) | | | | | | |
| Protein | 63.8 | 53.7 | 50.5 | 45.9 | 75.8 | 71.4 |
| Others | 95.4 | 75.3 | 82.9 | 75.1 | 104.9 | 109.4 |
| Waters | 47.4 | 48.7 | 46.7 | 27.7 | 52.8 | 48.0 |
| R.m.s. deviations | | | | | | |
| Bond lengths (Å) | 0.0037 | 0.0026 | 0.0037 | 0.0018 | 0.0019 | 0.0017 |
| Bond angles ($^O$) | 0.61 | 0.54 | 0.61 | 0.46 | 0.47 | 0.46 |
| Ramachandran plot (%) | | | | | | |
| Favored | 95.88 | 96.44 | 96.86 | 95.40 | 95.45 | 95.62 |
| Allowed | 4.03 | 3.38 | 3.00 | 4.42 | 4.37 | 4.07 |
| Outliers | 0.09 | 0.18 | 0.13 | 0.18 | 0.18 | 0.31 |
| **PDB ID** | 8OQQ | 8OQR | 8OQS | 8OQT | 8OQU | 8OQV |

[#] As reported in the PDB validation report

**Supplementary Table S4.** Summary of the fragment binding events at the subsites defined in **Table 3**.

| Fragment | occupancy | RSCC$^{\#}$ | ECH active site | | | HAD active site | | | | KAT active site | | | CoA-A$^{*}$ | CoA-B$^{*}$ | CoA-C$^{*}$ | | | other |
|---|---|---|---|---|---|---|---|---|---|---|---|---|---|---|---|---|---|---|
| | | | E1 | E2 | E3 | H0 | H1 | H2 | H3 | K1 | K2 | K3 | C1 | C2 | I1 | I2 | I3 | |
| B-E1 | | | | | | | | | | | | | | | | | | |
| O8D-B808 | 0.78 | 0.82 | | | | | | | | | | | | | | yes, B | | |
| O8D-B809 | 0.78 | 0.86 | | | | | | yes, B | | | | | | | | | | |
| O8D-D507 | 0.83 | 0.89 | | | | | | | | | | yes, D | | | | | | |
| O8D-C507 | 0.83 | 0.91 | | | | | | | | | | yes, C | | | | | | |
| O8D-A806 | 0.78 | 0.86 | | | | | | yes, A | | | | | | | | | | |
| B-H11 | | | | | | | | | | | | | | | | | | |
| STL-A806 | 1.00 | 0.88 | | | yes, A | | | | | | | | | | | | | |
| STL-A807 | 1.00 | 0.91 | | | | | | | | | | | | | yes, A | | | |
| STL-B806 | 1.00 | 0.86 | | | yes, B | | | | | | | | | | | | | |
| STL-B807 | 1.00 | 0.93 | | | | | | | | | | | | | yes, B | | | |
| B-51 | | | | | | | | | | | | | | | | | | |
| CFF-D507 | 1.00 | 0.92 | | | | | | | | | | | yes, D | | | | | |
| CFF-B810 | 1.00 | 0.93 | | | | | | | | | | | yes, C | | | | | |
| B-B3 | | | | | | | | | | | | | | | | | | |
| GLC-G1/G2 | 0.6/0.6 | 0.79/0.77 | | | | | | | | | | | | | | | | yes, D |
| GLC-F1/F2 | 0.8/0.8 | 0.86/0.78 | | | | | | | | | | | | | | | | yes, C |
| B-77 | | | | | | | | | | | | | | | | | | |
| DNQ-D501 | 0.5 | 0.77 | | | | | | | | | | | | | | | | yes, D |
| M-1 | | | | | | | | | | | | | | | | | | |
| A9J-A810 | 1.00 | 0.99 | | | yes, A | | | | | | | | | | | | | |
| A9J-A811 | 1.00 | 0.96 | | | | | | | | | | | | | | | | yes, A |
| A9J-A812 | 1.00 | 0.94 | | | | | | | | | | | | | | yes, A | | |
| A9J-A813 | 1.00 | 0.98 | | | | | | | | | | | | | | | | yes, A |
| A9J-A814 | 1.00 | 0.97 | | yes, A | | | | | | | | | | | | | | |
| A9J-A815 | 1.00 | 0.94 | | | | | | | | | | | | | | yes, A | | |
| A9J-A816 | 1.00 | 0.96 | | | | | | yes, A | | | | | | | | | | |
| A9J-A817 | 1.00 | 0.95 | | | | yes, A | | | | | | | | | | | | |
| A9J-A818 | 1.00 | 0.96 | | | | | | | | | | | | | | | | yes, A |
| A9J-A819 | 1.00 | 0.90 | | | | | | | | | | | | | | yes, A | | |
| A9J-B809 | 1.00 | 0.97 | | | yes, B | | | | | | | | | | | | | |
| A9J-B810 | 1.00 | 0.99 | | | | | | | | | | | | | | | | yes, B |
| A9J-B811 | 1.00 | 0.99 | | | | yes, B | | | | | | | | | | | | |
| A9J-B812 | 1.00 | 0.94 | | | | | | | | | | | | | | | | yes, B |
| A9J-B813 | 1.00 | 0.93 | | | | | | | | | | | | | | yes, B | | |

| | | | | | | | | | | | | |
|---|---|---|---|---|---|---|---|---|---|---|---|---|
| A9J-B814 | 1.00 | 0.98 | | yes, B | | | | | | | | |
| A9J-B815 | 1.00 | 0.95 | | | | | | | | yes, B | | |
| A9J-B816 | 1.00 | 0.91 | | | | | | | | yes, B | | |
| A9J-B817 | 1.00 | 0.96 | | | | yes, B | | | | | | |
| A9J-C510 | 1.00 | 0.99 | | | | | | | | | | yes, C |
| A9J-D508 | 1.00 | 0.98 | | | | | | yes, D | | | | |
| A9J-C511 | 1.00 | 0.98 | | | | | | yes, C | | | | |
| A9J-D509 | 1.00 | 0.99 | | | | | | | | | | yes, D |
| A9J-D510 | 1.00 | 0.90 | | | | | | | | | | yes, D |
| A9J-B818 | 1.00 | 0.89 | | | | | | | | | | yes, B |
| M-10 | | | | | | | | | | | | |
| VWO-A806 | 1.00 | 0.83 | yes, A | | | | | | | | | |
| VWO-B808 | 1.00 | 0.85 | yes, B | | | | | | | | | |
| VWO-D505 | 0,5 | 0.93 | | | | | | | | | | yes, D |
| M-49 | | | | | | | | | | | | |
| VXH-A2405 | 1.00 | 0.96 | | yes, A | | | | | | | | |
| VXH-A2406 | 1.00 | 0.91 | | | | | | | | yes, A | | |
| VXH-B2405 | 1.00 | 0.95 | | yes, B | | | | | | | | |
| VXH-B2406 | 1.00 | 0.94 | | | | | | | | | | yes, B |
| VXH-B2407 | 1.00 | 0.83 | | | yes, B | | | | | | | |
| VXH-2408 | 1.00 | 0.88 | | | | | | | | yes, B | | |
| VXH-C505 | 1.00 | 0.94 | | | | | | | | | | yes, C |
| VXH-D506 | 0.5/0.5 | 0.87/0.87 | | | | | | | | | | yes, D |
| M-53 | | | | | | | | | | | | |
| W3U-A805 | 1.00 | 0.91 | | | | | | | | | | yes, A |
| W3U-A806 | 1.00 | 0.86 | | | yes, A | | | | | | | |
| W3U-A807 | 0.5/0.5 | 0.82/0.82 | | | | | | | | | yes, A | |
| W3U-A808 | 1.00 | 0.77 | | | | | | | | | | yes, A |
| W3U-A809 | 1.00 | 0.89 | | | | | yes, A | | | | | |
| W3U-B809 | 1.00 | 0.83 | | | | | | | | yes, B | | |
| W3U-B810 | 0.5/0.5 | 0.83/0.83 | | | | | | | | | yes, B | |
| W3U-B811 | 0.5/0.5 | 0.86/0.86 | | | | | | | | | | yes, B |
| W3U-B812 | 1.00 | 0.87 | | | yes, B | | | | | | | |
| W3U-B813 | 1.00 | 0.91 | | | | | | | yes, B | | | |
| W3U-B814 | 1.00 | 0.92 | | | | | yes, B | | | | | |
| W3U-A810 | 1.00 | 0.72 | | | | | | | | yes, A | | |
| M-72 | | | | | | | | | | | | |
| JXL-B810 | 1.00 | 0.90 | | | | | | | | | yes, B | |
| JXL-B808 | 1.00 | 0.86 | yes, B | | | | | | | | | |

| | | | | | | | | | | | |
|---|---|---|---|---|---|---|---|---|---|---|---|
| YLZ-B805 | 1.00 | 0.93 | | | yes, B | | | | | | |
| JXL-B809 | 1.00 | 0.85 | | | | | | yes, B | | | |
| JXL-D508 | 1.00 | 0.86 | | | | | yes, D | | | | |
| JXL-C508 | 1.00 | 0.88 | | | | | yes, C | | | | |
| JXL-A810 | 1.00 | 0.92 | | | | | | | | yes, A | |
| JXL-A809 | 1.00 | 0.82 | yes, A | | | | | | | | |
| YLZ-A808 | 1.00 | 0.93 | | | yes, A | | | | | | |
| JXL-A811 | 1.00 | 0.83 | | | | | | yes, A | | | |
| YLN-A812 | 1.00 | 0.96 | | | yes, A | | | | | | |
| YMK-A806 | 1.00 | 0.92 | | | | | | | yes, A | | |
| YLZ-A807 | 1.00 | 0.95 | | | | | | | yes, A | | |
| YLN-B811 | 1.00 | 0.96 | | | yes, B | | | | | | |
| YLZ-B806 | 1.00 | 0.94 | | | | | | | yes, B | | |
| M-76 | | | | | | | | | | | |
| VXZ-A807 | 1.00 | 0.95 | | yes, A | | | | | | | |
| VXZ-A808 | 1.00 | 0.98 | | | | | | | | | yes, A |
| VXZ-A809 | 1.00 | 0.95 | | | | | | | | | yes, A |
| VXZ-A810 | 1.00 | 0.80 | | | | | | | | | yes, A |
| VXZ-A811 | 1.00 | 0.78 | | | | yes, A | | | | | |
| VXZ-B806 | 1.00 | 0.97 | | | | | | | | | yes, B |
| VXZ-B807 | 1.00 | 0.95 | | yes, B | | | | | | | |
| VXZ-B808 | 1.00 | 0.83 | | | | yes, B | | | | | |
| VXZ-B809 | 1.00 | 0.83 | | | | | | | | | yes, B |
| VXZ-B810 | 1.00 | 0.86 | | | | | | | | | yes, B |
| VXZ-B811 | 1.00 | 0.88 | | | | | | | | | yes, B |
| VXZ-C506 | 1.00 | 0.95 | | | | | | | | | yes, C |
| VXZ-C507 | 1.00 | 0.90 | | | | | | | | | yes, C |
| VXZ-D506 | 1.00 | 0.95 | | | | | | | | | yes, D |
| VXZ-D507 | 1.00 | 0.94 | | | | | | | | | yes, D |
| VXZ-D508 | 1.00 | 0.88 | | | | | | | | | yes, D |
| VXZ-A812 | 1.00 | 0.81 | | | | | | | | | yes, A |
| VXZ-D509 | 0.5 | 0.91 | | | | | | | | | yes, D |
| VXZ-B812 | 1.00 | 0.93 | | | | | | | | | yes, B |
| VXZ-A813 | 1.00 | 0.95 | | | | | | | | | yes, A |
| M-79 | | | | | | | | | | | |
| VWE-A805 | 1.00 | 0.95 | | yes, A | | | | | | | |
| VWE-A806 | 1.00 | 0.95 | | | | | | | | | yes, A |
| VWE-B808 | 1.00 | 0.92 | | yes, B | | | | | | | |
| VWE-B809 | 1.00 | 0.95 | | | | | | | | | yes, B |

| | | | | | | | | | | | | | | | | | |
|---|---|---|---|---|---|---|---|---|---|---|---|---|---|---|---|---|---|
| M-80 | | | | | | | | | | | | | | | | | |
| VWT-A808 | 1.00 | 0.89 | | yes, A | | | | | | | | | | | | | |
| VWT-A809 | 1.00 | 0.88 | yes, A | | | | | | | | | | | | | | |
| VWT-B807 | 1.00 | 0.84 | | yes, B | | | | | | | | | | | | | |
| VWT-B808 | 1.00 | 0.95 | yes, B | | | | | | | | | | | | | | |
| VWT-A810 | 1.00 | 0.95 | | | yes, B | | | | | | | | | | | | |
| VWT-A811 | 1.00 | 0.98 | | | yes, A | | | | | | | | | | | | |
| M-83 | | | | | | | | | | | | | | | | | |
| VWZ-A809 | 1.00 | 0.95 | | | | | | yes, A | | | | | | | | | |
| VWZ-A810 | 0.5/0.5 | 0.85/0.85 | | | | yes, A | | | | | | | | | | | |
| VWZ-A811 | 1.00 | 0.86 | | yes, A | | | | | | | | | | | | | |
| VWZ-A812 | 1.00 | 0.92 | | yes, A | | | | | | | | | | | | | |
| VWZ-A813 | 1.00 | 0.79 | yes, A | | | | | | | | | | | | | | |
| VWZ-A814 | 1.00 | 0.83 | | | | | yes, A | | | | | | | | | | |
| VWZ-B810 | 0.5/0.5 | 0.91/0.91 | | | | yes, B | | | | | | | | | | | |
| VWZ-B811 | 1.00 | 0.87 | | yes, B | | | | | | | | | | | | | |
| VWZ-B812 | 1.00 | 0.90 | | yes, B | | | | | | | | | | | | | |
| VWZ-B813 | 1.00 | 0.74 | yes, B | | | | | | | | | | | | | | |
| VWZ-B814 | 1.00 | 0.81 | | | | | yes, B | | | | | | | | | | |
| VWZ-C508 | 1.00 | 0.89 | | | | | | | | | yes, C | | | | | | |
| VWZ-D508 | 1.00 | 0.91 | | | | | | | | | yes, D | | | | | | |
| VWZ-B815 | 1.00 | 0.89 | | | | | | | | | | | | | | | yes, B |
| VWZ-A815 | 1.00 | 0.91 | | | | | | | | | | | | | | | yes, A |
| VWZ-B816 | 1.00 | 0.90 | | | | | | yes, B | | | | | | | | | |
| M-91 | | | | | | | | | | | | | | | | | |
| VXC-A808 | 1.00 | 0.91 | yes, A | | | | | | | | | | | | | | |
| VXC-A809 | 1.00 | 0.90 | | yes, A | | | | | | | | | | | | | |
| VXC-B809 | 1.00 | 0.92 | yes, B | | | | | | | | | | | | | | |
| VXC-B810 | 1.00 | 0.91 | | yes, B | | | | | | | | | | | | | |
| VXC-B811 | 1.00 | 0.80 | | yes, B | | | | | | | | | | | | | |
| VXC-A810 | 1.00 | 0.96 | | yes, A | | | | | | | | | | | | | |
| M-92 | | | | | | | | | | | | | | | | | |
| VXN-B807 | 1.00 | 0.94 | | yes, B | | | | | | | | | | | | | |
| VXN-B808 | 1.00 | 0.89 | | yes, B | | | | | | | | | | | | | |
| VXN-A808 | 1.00 | 0.96 | | yes, A | | | | | | | | | | | | | |
| VXN-B809 | 1.00 | 0.81 | yes, B | | | | | | | | | | | | | | |
| VXN-A809 | 1.00 | 0.91 | | yes, A | | | | | | | | | | | | | |
| VXN-A810 | 1.00 | 0.87 | yes, A | | | | | | | | | | | | | | |
| M-109 | | | | | | | | | | | | | | | | | |

| | | | | | | | | | | | | | | | | | | |
|---|---|---|---|---|---|---|---|---|---|---|---|---|---|---|---|---|---|---|
| VWI-A806 | 1.00 | 0.89 | | | | | | | yes, A | | | | | | | | | |
| VWI-A807 | 1.00 | 0.78 | | | | | | yes, A | | | | | | | | | | |
| VWI-A808 | 1.00 | 0.90 | | | yes, A | | | | | | | | | | | | | |
| VWI-B804 | 1.00 | 0.93 | | | yes, B | | | | | | | | | | | | | |
| VWI-A809 | 1.00 | 0.76 | | yes, A | | | | | | | | | | | | | | |
| VWI-B805 | 1.00 | 0.78 | | yes, B | | | | | | | | | | | | | | |
| VWI-A810 | 1.00 | 0.86 | | | yes, A | | | | | | | | | | | | | |
| VWI-B806 | 1.00 | 0.85 | | | | | | | yes, B | | | | | | | | | |
| | | | E1 | E2 | E3 | H0 | H1 | H2 | H3 | K1 | K2 | K3 | C1 | C2 | I1 | I2 | I3 | other |
| Totals (143) | | | 4 | 18 | 28 | 4 | 4 | 7 | 6 | 0 | 0 | 8 | 2 | 3 | 3 | 13 | 4 | 39 |
| | | | | ECH | | | HAD | | | | KAT | | CoA-A* | CoA-B* | | CoA-C* | | |
| Totals (143) | | | | 50 | | | 21 | | | | 8 | | 2 | 3 | | 20 | | 39 |

[#] As reported in the PDB validation report.

* CoA-A, CoA-B, CoA-C refer to the CoA-A(HAD/KAT), CoA-B(ECH2) and CoA-C(ECH/HAD) regions, respectively.

**Supplementary Table S5.** The 15 fragment binding subsites and the fragments that are bound at these subsites.[a]

| Binding site | Subsite | Subsite defining residues | Fragment (number of binding events) |
|---|---|---|---|
| ECH active site | E1 | αGly67, αLeu114, αGly115, αPro140, αLeu144, αArg175, αPhe304 | M-10(2), M-72(2) |
| | E2 | αGly67, αGly68, αAsp69, αVal70, αMet73, αThr143, αLeu114, αGly115, αGly116, αPro140, αGlu141, αThr143, αLeu144, αArg175, αPhe304 | M-1(2), M-49(2), M-76(2), M-79(2), M-80(2), M-83(2), M-91(2), M-92(2), M-109(2) |
| | E3 | αMet30, αAsn31, αGlu32, αIle35, αGly68, αAsp69, αThr72, αMet73, αAla76, αAsp80, αGlu82, αAsp83, αVal84, αThr87, αIle91, αGly116, αGlu119, αGlu141, αGly149, αGly150, αPhe287 | B-H11(2), M-1(2), M-49(1), M-53(2), M-72(4), M-80(2), M-83(4), M-91(4), M-92(4), M-109(3) |
| HAD active site | H0 | αLeu331, αGly332, αLys354, αAsp355, αVal356, αAla411, αVal412, αVal412, αPhe413, αAsn439, αMet470, αPro471, αPro545 | M-1(2), M-80(2) |
| | H1 | αMet470, αPro471, αIle667, αMet668, αIle670, αGly671, αPro545 | M-76(2), M83 (2) |
| | H2 | αSer441, αHis462, αPhe464, αGly508, αSer512, αIle515, αArg513, αAsn520, αLeu555, αLeu559, αMet560, αI563, αGly669, αIle670 | B-E1(2), M-1(2), M-83(2), M-109(1) |
| | H3 | αThr442, αPro444, αArg507, αGly508,αSer512, αArg513, αGly516, αVal519, αAsn520, αArg507, αLeu559, αMet560, αIle563, αAla566, αThr567 | M-53(2), M-83(2), M-109(2) |
| KAT active site | K1 | | no bound fragments |
| | K2 | | no bound fragments |
| | K3 | βGly67, βArg71, βAla72, βLeu75, βPhe91, βVal147, βMet134, βPhe146, βGln149, βPro296, βMet299, βGly391, βGly392 | B-E1(2), M-1(2), M-72(2), M-83(2) |
| CoA-C(ECH/HAD) site | I1 | αLeu14, αThr43, αAla66, αGly67, αLeu114, αPro140, αGlu141, αThr143, αThr143, αLeu144, αArg175, αPhe303, αLeu307, | B-H11(2), M-72(1) |
| | I2 | αThr143, αIle167, αPhe166, αAla171, αGln172, αLeu249, αGln252, αLeu253, αMet258, αPhe303, αMet470, αPro471, αPro545, αIle667, αMet668 | B-E1(1), M-1(6), M-49(2), M-53(2), M-72(2) |
| | I3 | αAla161, αPhe166, αGln163, αVal167, αIle171, αIle241, αSer244, αPhe245, αAsn248, αLeu249, αGln252, βLeu231 | M-53(2), M-72(2) |
| CoA-A(HAD/KAT) site | C1 | αGln629, βTrp244 | B-51(2) |
| CoA-B(ECH2) site | C2 | αPhe160, αAsn164, αSer168, αLys180, αIle184 | M-53(1), M-72(2) |

[a]The listed residues have atoms within 3.7Å of the bound fragment. It concerns 104 binding events, only two binding events (of fragment M-80 at subsite H0) are at a crystal contact.

**Supplementary Table S6.** Analysis of the CoA-protein interactions of CoA bound in the active sites and in the additional binding sites.

| **Active sites** | **PDB ID/ residue number** | **SC/ total** | **SC perentage** | **S-S interactions** | **Salt bridge interactions** | **H-bond/ total** | **H-bond percentage** | **Interactions of the adenine moiety** |
|---|---|---|---|---|---|---|---|---|
| CoA (ECH) - chain A (bent conformation) | 7O4T/A801 | 15/23 | | 0 | 0 | 9/23 | | Stacking of adenine between side chains of αPhe304 and αVal29 |
| CoA (ECH) - chain B (bent conformation) | 7O4T/B801 | 20/30 | | 0 | 0 | 12/30 | | |
| CoA (KAT) - chain C (extended conformation) | 7O4T/C501 | 19/24 | | 1 | 0 | 6/24 | | Stacking of adenine between side chains of βArg210 and βLeu221 |
| CoA (KAT) - chain D (extended conformation) | 7O4T/D501 | 26/33 | | 1 | 1 (weak) | 7/33 | | |
| CoA active sites, totals: | | | | | | | | |
| CoA (total) | | 80/110 | 73% | | | 34/110 | 31% | |
| CoA (average) | | 20.0/27.5 | 73% | | | 8.5/27.5 | 31% | |
| **Additional binding sites** | | | | | | | | |
| CoA-A(HAD/KAT) chains BC | 7O4R/D506 | 30/31 | | 0 | 1 | 4/31 | | Stacking of adenine between side chains of βTrp244 and αGln629 |
| CoA-A(HAD/KAT) chains AD | 7O4R/C508 | 25/25 | | 0 | 1 | 3/25 | | |
| CoA-B(ECH2) chain A | 7O4S(A809 | 23/27 | | 0 | 1 | 5/27 | | Stacking of adenine between pantetheine moiety and bulk solvent |
| CoA-B(ECH2) chain B | 7O4S/B808 | 18/25 | | 0 | 1 | 6/25 | | |
| CoA-C(ECH/HAD) chain A | 7O4T/A802 | 14/24 | | 0 | 0 | 1/24 | | Stacking of adenine between side chain of αLys469 and bulk solvent |
| CoA-C(ECH/HAD) chain B | 7O4T/B803 | 5/8 | | 0 | 0 | 0/8 | | |
| CoA additional binding sites, totals: | | | | | | | | |
| CoA (total) | | 115/140 | 82% | | | 19/140 | 14% | |
| CoA (average) | | 19.2/23.3 | 82% | | | 3.2/23.3 | 14% | |

The contact distance cut-off value has been 3.7Å. None of the additional CoA binding sites are near crystal contacts. SC refers to interactions with side chains. H-bond refers to hydrogen bond interactions. S-S-interactions refer to interactions between two sulfur atoms. These S-S interactions are only observed in the thiolase active site and they have been classified as Van der Waals interactions (in both active sites of the thiolase dimer, it concerns interactions with the sulfur atom of the nucleophilic cysteine (at 3.2Å); the other, nearest sulfur is from Cys389, the acid/base cysteine, which is at 5Å). Hydrogen bond interactions are counted after visual inspection.

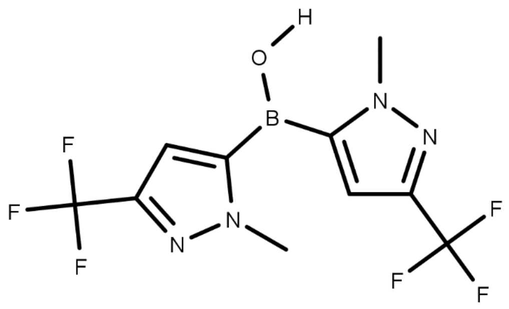


**Figure S1**. The covalent structure of the M-72 dimer.

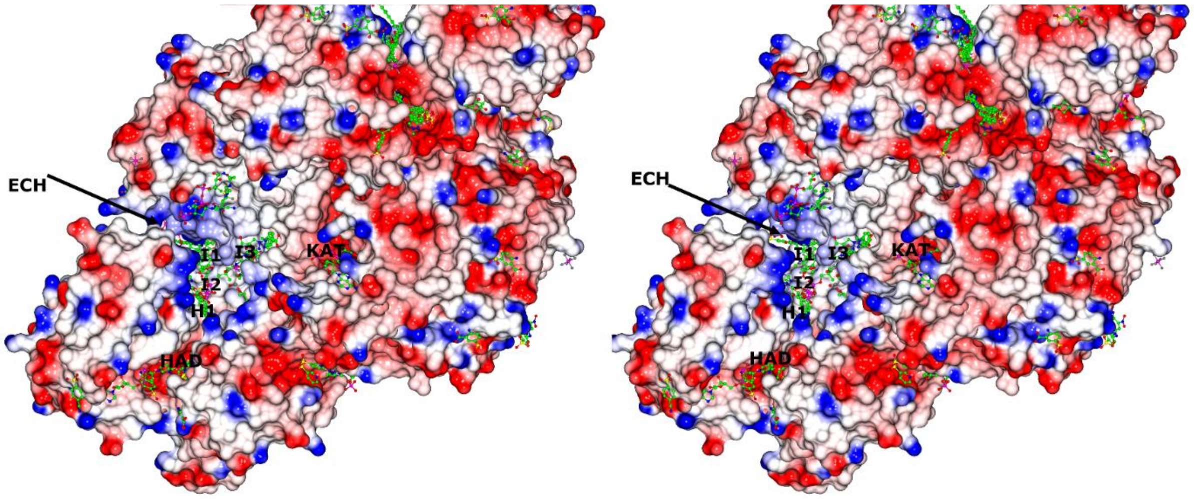


**Figure S2**. Stereo image of the protein surface of the possible substrate channeling path between the three active sites of MtTFE, as also shown in **Fig. 5**. The image shows the electrostatic properties of the surface of the CoA-C(ECH/HAD) region, near the interface of chain A (α subunit) and chain D (β subunit) of the CoA-C structure (PDB ID 7O4T), color coded such that the blue and red colors identify surface regions with positive and negative electrostatic potential, respectively. The locations of the catalytic sites are labeled as ECH (with arrow), HAD and KAT. The neutral (white color) CoA-C(ECH/HAD) site is labeled as I2, whereas the regions extending to the ECH, HAD and KAT catalytic sites are labeled as I1, H1, and I3.

**Movie S1.** The fragment binding events at the surface region near the CoA-C(ECH//HAD) binding site between the ECH, HAD and KAT active sites. The electrostatic surface is calculated from the CoA-C structure (PDP ID 7O4T), color coded such that the blue and red colors identify surface regions with positive and negative electrostatic potential, respectively. In the very first frames the MtTFE cartoon structure is shown, together with ligands bound in the ECH active site (CoA), HAD active site ($NAD^+$) and KAT active site (CoA) as well as the CoA bound in the three additional CoA binding sites. All 143 fragment binding events are included in the subsequent frames. The view is similar as in **Fig. 1** and **Fig. 5**.
